# Surprising amount of stasis in repetitive genome content across the Brassicales

**DOI:** 10.1101/2020.06.15.153296

**Authors:** Aleksandra Beric, Makenzie E. Mabry, Alex E. Harkess, M. Eric Schranz, Gavin C. Conant, Patrick P. Edger, Blake C. Meyers, J. Chris Pires

**Affiliations:** Donald Danforth Plant Science Center, St. Louis, Missouri 63132, Division of Plant Sciences, University of Missouri, Columbia, Missouri 65211; Division of Biological Sciences and Bond Life Sciences Center, University of Missouri, Columbia, Missouri 65211; Auburn University, Department of Crop, Soil, and Environmental Sciences, Auburn, AL 36849, HudsonAlpha Institute for Biotechnology, Huntsville, Alabama 35806; Wageningen University and Research, Wageningen, Netherlands; Bioinformatics Research Center, Program in Genetics and Department of Biological Sciences, North Carolina State University, Raleigh, North Carolina 27695; Department of Horticulture, Department of Ecology, Evolutionary Biology and Behavior, Michigan State University, East Lansing, Michigan 48824

**Keywords:** Brassicales, Repetitive Elements, Whole-genome duplication, Genome size, Evolution

## Abstract

Genome size of plants has long piqued the interest of researchers due to the vast differences among organisms. However, the mechanisms that drive size differences have yet to be fully understood. Two important contributing factors to genome size are expansions of repetitive elements, such as transposable elements (TEs), and whole-genome duplications (WGD). Although studies have found correlations between genome size and both TE abundance and polyploidy, these studies typically test for these patterns within a genus or species. The plant order Brassicales provides an excellent system to test if genome size evolution patterns are consistent across larger time scales, as there are numerous WGDs. This order is also home to one of the smallest plant genomes, *Arabidopsis thaliana* - chosen as the model plant system for this reason - as well as to species with very large genomes. With new methods that allow for TE characterization from low-coverage genome shotgun data and 71 taxa across the Brassicales, we find no correlation between genome size and TE content, and more surprisingly we identify no significant changes to TE landscape following WGD.

## INTRODUCTION

Genome sizes across flowering plants (angiosperms) vary from 65 Mbp in Lentibulariaceae (Fleischmann *et al.* 2014), a family of carnivorous plants, to approximately 150 Gbp in *Paris japonica* (Pellicer *et al.* 2010), making it not only the largest genome in the angiosperms but also within all Eukaryotes (Hidalgo *et al.* 2017). In the Brassicales, an economically important order of plants in the angiosperms, genome sizes range from 156 Mbp to 4.639 Gbp, with both extremes coming from the Brassicaceae family *(Arabidopsis thaliana;* Bennett *et al.* 2003 and *Crambe cordifolia;* Lysak *et al.* 2007). This incredible breadth in genome size among plant species cannot be explained solely by the number of protein-coding genes, a discrepancy known as the “C-value paradox” (Thomas, 1971). Instead, genome size and its evolution are largely influenced by the amount of non-coding sequences and repetitive elements (Elliott and Gregory, 2015). There are several theories trying to explain what gave rise to this paradox. Some suggest that lack of natural selection, possibly due to small effective population sizes, allowed accumulation of DNA material that would otherwise get purged from the genome (Lynch and Conery 2003; Doolittle 2013). Others postulate that non-coding DNA was selectively expanded to enable increase in cell size, thus lowering the metabolic rate, and permitting overall increase in body size at a lower cost (Kozlowski *et al.* 2003). Indeed, the question of the factors driving genome size connects to deep questions regarding the structure of genomes, the interplay of the natural selection and population size, and even the relationship between body size to metabolic rate.

Large portions of plant genomes are made up of a particular type of repetitive elements, known as transposable elements (TEs). These are mobile repetitive elements which are dispersed throughout the genome (Kubis *et al.* 1998). TEs are grouped in two major classes, based on their mechanism of transposition. Each of the two classes is further resolved into superfamilies, which vary in repeat domain structure. Class I TEs (or retrotransposons) move to a new genomic location via an RNA intermediate, a mechanism commonly called “copy-paste” (Wicker *et al.* 2007; Negi *et al.* 2016). This copy-paste mechanism results in an increased copy number of a retrotransposon (Wicker *et al.* 2007). Retrotransposons code for a reverse transcriptase, which is a defining component of their transposition mechanism. Plant genomes are often dominated by the two high-copy Class I TE superfamilies: *Copia* and *Gypsy* (Macas *et al.* 2015; Wicker *et al.* 2018).

Class II TEs (or DNA transposons) are defined by a “cut-paste” mechanism of transposition, which utilizes a DNA intermediate. The majority of Class II elements are characterized by two main features: terminal inverted repeats (TIRs) and a transposase enzyme (Wicker *et al.* 2007; Negi *et al.* 2016). One such superfamily is *Mutator.* In order for *Mutator* elements to move from one genomic location to another, a transposase needs to first recognize the TIRs and then cut both DNA strands on either end of the TE (Wicker *et al.* 2007). Insertion of the TE into a new location results in a small target site duplication (TSD). A TSD is a signature typical of DNA element transposition, as the target site will remain duplicated when the TE is excised and moves to another location in the genome (Muñoz-López and García-Pérez, 2010). Although they typically move in a cut-paste fashion, transposition of Class II elements can lead to an increase in their copy number when they are inserted in front of a replication fork (Wicker *et al.* 2007).

The effects of TE movement are not necessarily deleterious and have been shown to cause maize kernel variegation (McClintock, 1950). TE insertions have also been found to be the source of other economically important phenotypic variation such as grape berry color, morning glory flower variegation, and parthenocarpic apple fruit (Kobayashi *et al.* 2004; Cadle-Davidson and Owens, 2008; Shimazaki *et al.* 2011; Bennetzen, 2005; Clegg and Durbin, 2000; Habu *et al.* 1998; Yao *et al.* 2001). All of these effects point to TEs as large contributors to genome evolution and plasticity.

The abundances of TEs in genomes have also been shown to be informative when inferring phylogenetic relationships among taxa, especially in groups rife with polyploidy, such as those in the Brassicales, because it adds an analysis complementary to species tree phylogenies (Dodsworth *et al.* 2015; Harkess *et al.* 2016; Dodsworth *et al.* 2017; Vitales *et al.* 2020). Several studies have used maximum parsimony methods to reconstruct phylogenetic trees, treating TE abundances as continuous characters (Dodsworth *et al.* 2015; Dodsworth *et al.* 2017). The resulting trees are largely concordant to those produced via traditional phylogenetic methods (i.e., using protein-coding genes). More recently, a study has shown the power of combining TE sequence similarity with TE abundance to understand evolutionary relationships (Vitales *et al.* 2020). Generally, *Copia* and *Gypsy* are the most informative elements due to their high abundance in the genomes, whereas low-abundant TEs are insufficient to resolve the phylogenetic relations with these approaches (Dodsworth *et al.* 2015; Harkess *et al.* 2016; Dodsworth *et al.* 2017; Vitales *et al.* 2020).

Polyploidy or whole-genome duplication (WGD) is typically followed by extensive chromosomal rearrangements, gene loss, and epigenetic remodeling during the process of diploidization (Schranz and Mitchell-Olds 2006, Madlung *et al.* 2005). This genome restructuring has been associated with both expansion and loss of TEs (Ågren *et al.* 2016; Parisod *et al.* 2010; Vicient and Casacuberta, 2017). TE mobilization and proliferation following WGD have been recorded in tobacco, wheat, as well as many Brassicaceae species (Ben-David *et al.* 2013; Petit *et al.* 2010; Sarilar *et al.* 2011; Ågren *et al.* 2016; Vicient and Casacuberta, 2017). TE amplification in wheat, however, seems to be family specific, as there is no evidence of massive reactivation of TEs (Wicker *et al.* 2018). In fact, TE abundance, landscape in gene vicinity, and the proportion of different TE families show surprising levels of similarity between the three wheat subgenomes. While there is evidence of large TE turnovers after the divergence of the A, B, and D subgenomes, these turnovers seem to have happened prior to hybridization (Wicker *et al.* 2018). Following polyploidization, TEs can accumulate in regions proximal to genes and gene-regulatory elements, leading to dynamic variation in gene expression (Ågren *et al.* 2016; Sarilar *et al.* 2011; Negi *et al.* 2016).

The Brassicales is a particularly valuable order as a model in which to elucidate the connection between whole genome duplication and repetitive element proliferation. First, there are at least four major polyploid events in the Brassicales: “At – α” at the base of the Brassicaceae, the Brassiceae whole-genome triplication (“WGT”), “Th – α” in the Cleomaceae (Schranz and Mitchell-Olds 2006; Barker *et al.* 2009; Mabry *et al.* 2020), and “At – β” at the base of the order (Edger *et al.* 2015; 2018). Second, genomes within the Brassicales are typically small, less than 500 Mb. Several genome projects have produced highly contiguous genome assemblies with accurate gene and repeat annotations, such as *Arabidopsis thaliana* (The Arabidopsis Genome Initiative 2000; Michael *et al.* 2018), several *Brassica* sp. genomes (Wang *et al.* 2011; Liu *et al.* 2014; Parkin *et al.* 2014; Chalhoub et al. 2014), *Cleome violacea* (Emery *et al.* 2018), *Carica papaya* (Nagarajan *et al.* 2008), and *Thlaspi arvense* (Dorn *et al.* 2015).

Complex dynamics of TEs have been studied for a variety of species, but most studies focus on one or a few closely-related species. More robust comparative studies are vital for understanding large-scale patterns of TE behavior in response to evolutionary pressures. Highly contiguous, whole-genome assemblies and annotations are the gold standard for repetitive element annotation and their quantification within a genome, but this is a cost-prohibitive approach as the sample number increases. Several approaches, such as RepeatExplorer (Novak *et al.* 2013) and Transposome (Staton and Burke 2015), have been developed to use low-cost Illumina genome shotgun data to assess repetitive element content and abundance. Here, we leverage low coverage, genome shotgun data to examine both the relationship between 1) TE abundance and genome size, and 2) TE dynamics and WGD, in a dataset consisting of 71 taxa across the Brassicales.

## MATERIALS AND METHODS

### Taxon sampling, RNA and DNA isolation, and sequencing

Sampling of 71 taxa across the Brassicales spanned seven families and 57 genera, with a focus on the families Brassicaceae (47 taxa) and Cleomaceae (15 taxa; **Table S1**). Seeds were grown at the University of Missouri – Columbia or the University of Alberta in a sterile growth chamber environment. Leaf tissue from mature plants was collected for both RNA and DNA extraction followed by isolation and sequencing as in Mabry *et al.* (2020).

### Genome sizing

For estimation of genome size by flow cytometry, single leaves from 69 taxa were cut and placed in a wet paper towel and shipped to the Benaroya Research Institute at Virginia Mason (Seattle, WA). Nuclei isolations from a single mature leaf were analyzed in four technical replicates for each sample. Analyses were carried out using the Partec PAS flow cytometer (Partec, http://www.partec.de/), equipped with a mercury lamp. Leaves (0.1 g) were chopped in a nuclei extraction buffer (CyStain ultraviolet Precise P Nuclei Extraction Buffer; Partec, Münster, Germany) and filtered through a 30 mm Cell-Trics disposable filter (Partec), followed addition of 1.2 ml of staining solution containing 4,6-diamidino-2-phenylindole. The relative fluorescence intensity of stained nuclei was measured on a linear scale, and 4000 to 5000 nuclei for each sample were analyzed (Galbraith *et al.* 1998).

### Repetitive element analysis

Prior to repeat identification, DNA sequencing reads were paired to corresponding paired-end sequence files using a stand-alone Pairfq script, version 0.16.0 (https://github.com/sestaton/Pairfq) followed by removal of any reads that matched custom databases of mitochondrial and chloroplast genomes, comprised of sequences downloaded from NCBI (**Table S2**). Subsequently, low quality and short reads were removed with Trimmomatic v 0.39, with MINLEN of 70 and LEADING and TRAILING thresholds set to 3 (Bolger *et al.* 2014). Repeats were identified using Transposome v 0.12.1 (Staton and Burke 2015), which uses graph-based clustering, with 90% identity and fraction coverage of 0.55. Transposome was chosen over other similar software due to its speed, accessibility (i.e., open source), and ability to annotate repetitive elements from sequence data without the need for an assembled genome. For all but five species, repeat identification was performed with 500,000 random read pairs (see **Table S3** for details). These five samples had to be further downsampled due to script limitations (*see* How to choose the appropriate genome coverage section; https://github.com/sestaton/Transposome/wiki/). A database containing all repetitive elements previously annotated in the Viridiplantae was obtained from RepBase v 21.10 and used as reference for Transposome cluster annotation. The reported genomic fraction of each TE family represents the abundance of that family that has been corrected for by the number of unclustered reads (*see* Specifications and example usage section; https://github.com/sestaton/Transposome/wiki/).

### Correlation between genome size and TE content

The *pic* function from the ape package was used to calculate phylogenetically independent contrasts (PIC) (Paradis and Schliep 2018). The *lm* function from the stats package in R (version 3.6.1) was then used to perform linear regression analysis using the computed PIC values (R Core Team 2019). Correlation between genome size and total TE abundance, *Gyspy* abundance, and *Copia* abundance was studied on a subset of data, excluding species with genomes larger than 1.3Gbp *(Cakile maritima, Farsetia aegyptia, Hesperis matronalis, Matthiola longipetala, Physaria acutifolia, Schizopetalum walkeri).* These genomes were removed to avoid the bias caused by a small group of very large genomes present in Brassicaceae. Further, two other species were not included in these analyses, *Cleomella serrulata* due to the lack of DNA sequence and genome size data for this species, and *Sisymbrium brassiformis* due to lack of genome size data.

### Transcriptomics, phylogeny estimation, and hierarchical clustering

Transcriptome assembly and alignment follow Mabry *et al.* (2020), but in brief, reads were quality filtered and adapter-trimmed using Trimmomatic v 0.35 (Bolger *et al.* 2014) and assembled using Trinity v 2.2 (Grabherr *et al.* 2011) followed by translation to protein sequences using TransDecoder v 3.0 (github.com/TransDecoder/TransDecoder). Orthology was determermined using OrthoFinder v.2.2.6 (Emms and Kelly 2018) followed by filtering for taxon occupancy and alignment quality (github.com/MU-IRCF/filter_by_ortho_group, github.com/MU-IRCF/filter_by_gap_fraction). Gene trees were estimated using RAxML v 8 (Stamatakis 2014) followed by PhyloTreePruner v 1.0 (Kocot *et al.* 2013) to remove any potentially remaining paralogous genes. Final alignments were again run through RAxML v 8 (Stamatakis 2014), followed by species tree estimation using ASTRAL-III v.5.6.1 (Zhang *et al.* 2018).

To test for phylogenetic signal in repetitive element abundance, hierarchical clustering using *dist* and *hclust* functions from the stats package in R was conducted (R Core Team 2019). To construct a consensus dendrogram, hierarchical clusters produced with *Copia*, *Gypsy*, and total TE abundances were used. The consensus dendrogram was calculated using the mergeTree v.0.1.3 package in R (Hulot *et al.* 2019). *Cleomella serrulata* was not included in these analyses, due to the lack of DNA sequencing data for this species.

### Comparative genomics

To produce an ultrametric tree necessary for comparative genomic analyses, final alignments were concatenated (those without paralogous genes) using scripts from the Washburn *et al.* (2017) Genome-Guided Phylo-Transcriptomic pipeline (‘concatenate_matrices.py’), followed by tree estimation in RAxML v8 (Stamatakis 2014) with 100 bootstrap replicates and *Moringa* and *Carica* as outgroups. Branch lengths and model parameters were optimized using the ASTRAL phylogeny as a fixed input tree. Dating of the resulting tree was calculated in TreePL v. 1.0 (Smith and O’Meara 2012) using two fossils. *Palaeocleome lakensis* was used to date the node between the Cleomaceae and Brassicaceae (minimum age = 47.8, 95% highest posterior density = 52.58; Cardinal-McTeague, Sytsma, and Hall 2016); *Dressiantha bicarpellata* was used to date the node between the Caricaceae and Moringaceae, and the remaining Brassicales (minimum age = 89.9, 95% highest posterior density = 98.78; Cardinal-McTeague, Sytsma, and Hall 2016).

To test for the placement and magnitude of possible adaptive shifts in the data, Bayou v 2.0 was used (https://github.com/uyedaj/bayou/blob/master/tutorial.md; Uyeda and Harmon, 2014). *Cleomella serrulata* was again dropped from analyses due to lack of DNA sequence data for this sample. Analyses were run on total TE, *Gyspy*, and *Copia* abundances. Priors for all three analyses were as follows: lognormal distributions were used for alpha (the strength of the pull toward trait optima) and sigma squared (rate of phenotypic evolution) both with default parameters, a conditional Poisson distribution for the number of shifts using default parameters, a normal distribution for theta (the value of the optima) with mean equal to the mean of the observed data and standard deviation equal to two times the standard deviation of the data, and a uniform distribution for branch shifts with default parameters. Analyses were run for 1,000,000 generations and then checked for lack of convergence with a burn in of 0.3. Heatmaps of reconstructed values were plotted on tree branches using the *plotBranchHeatMap* function in Bayou. For Simmap trees, a posterior probability of 0.3 was used as a cutoff for shift identification.

To test the likelihood of WGDs being associated with identified shifts in TE abundances, OUwie v 2.1 was used (www.rdocumentation.org/packages/OUwie/; Beaulieu *et al.* 2012). Four separate selective regimes were used: 1) At – α at the base of the Brassicaceae versus all other Brassicales, 2) the tribe Brassiceae WGT versus all other Brassicaceae, 3) Th – α versus all other Cleomaceae, and 4) a reduced subset of 19 taxa for which we were confident in determining ploidy level, for which we tested diploids vs polyploids (**Table S1**). All seven models (single-rate Brownian motion; BM1, Brownian motion with different rate parameters for each state on a tree; BMS, Ornstein-Uhlenbeck model with a single optimum for all species; OU1, Ornstein-Uhlenbeck model with different state means and a single alpha and sigma^2 acting all selective regimes; OUM, Ornstein-Uhlenbeck model that assumes different state means as well as multiple sigma^2; OUMV, Ornstein-Uhlenbeck model that assumes different state means as well as multiple alpha; OUMA, Ornstein-Uhlenbeck model that assumes different state means as well as multiple alpha and sigma^2 per selective regime; OUMVA) were run for Total TE, *Gyspy*, and *Copia* abundances and compared using the weighted Akaike information criterion corrected for samplesize (AICc).

### Data availability

The authors state that all data necessary for confirming the conclusions presented in the article are represented fully within the article. Both RNA and DNA sequence data from this article can be found in the NCBI SRA data libraries under BioProject accession number PRJNA542714. Scripts are available at https://github.com/mmabry/Brassicales_RepetitiveElements.

## RESULTS AND DISCUSSION

### Sequence matrices and genome size patterns across the Brassicales

In order to investigate and interpret the evolution of genome size across the Brasicales, we first needed to construct a reliable phylogenetic framework based on our transcriptome assemblies. After determining orthology using OrthoFinder2, we recovered 35,522 orthogroups, with a final 1,404 orthogroups remaining after filtering. The resulting phylogeny had all but eight nodes recovered with a local posterior probability of 0.7 or greater (**Figure 1**). Next, repetitive element clustering and quantification using Transposome was performed for each species and mapped onto the species tree. After clustering, less than 2.1% of reads remain unannotated for any given species, likely reflective of the high quality of Brassicales genome annotations in RepBase (see **Table S3** for details). Total repeat content of species across the Brassicales ranged from 35.5% to 72.5% (mean 52.7, SD +/− 9.35; **Figure 1**). Overall, DNA-type “cut and paste” transposons comprise between 0.6% and 14.8% of genomes, largely dominated by *MuDR* and *Helitron* elements (**Figure 1**). In contrast, long terminal repeat (LTR) retrotransposons, mostly composed of *Ty3-Gypsy* and *Ty1-Copia* elements, tend to make up most of the total repeat content (**Figure S1**). Genome sizes ranged from 195.6 Mbp in *Descurainia sophioides* to 3261.63 Mbp in *Hesperis matronalis.* For two samples, *Cleomella serrulata* and *Sisymbrium brassiformis,* we were unable to obtain leaf tissue for genome sizing. The genome sizes recovered represent well the diversity and range of genome size that is found across the Brassicales (https://cvalues.science.kew.org/; Release 7.1, Leitch *et al.* 2019). The median and mean for families which were represented by more than one sample were as follows: Brassicaceae – 557.5 Mbp, 735.4 Mbp; Capparaceae – 511.0 Mbp, 605.1 Mbp; Cleomaceae – 479.2 Mbp, 549.1 Mbp, and Resedaceae – 684.6 Mbp, 684.6 Mbp (**Figure 1**).

**Figure 1.**
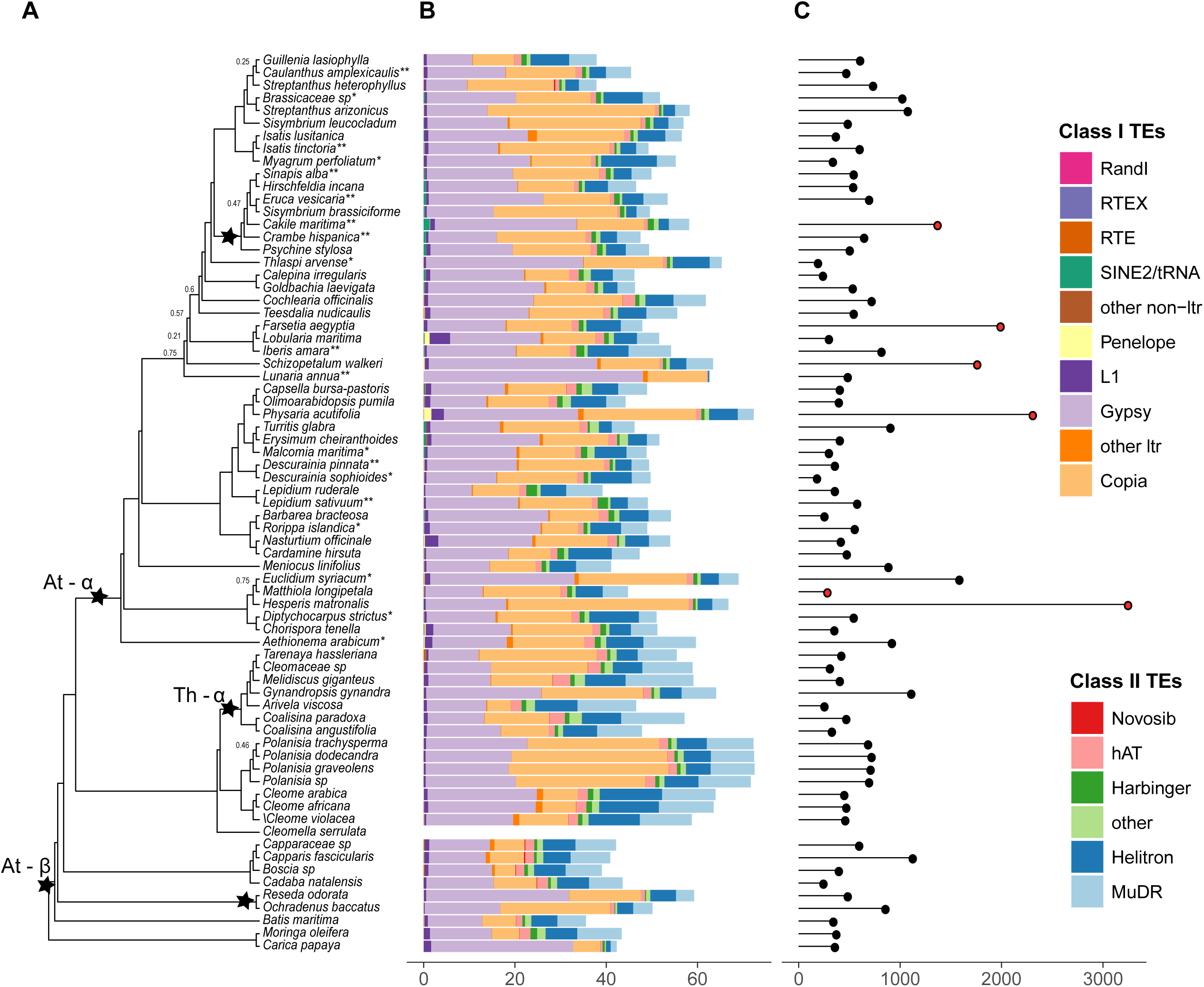
Taxa sampled with corresponding transposable element (TE) abundance and genome size. (A) Phylogeny with whole-genome duplications indicated by black stars. Support values are shown for those branched with less than 0.7 local posterior probability. Asterisks (*) next to taxon indicate known samples with known ploidy levels (** = polyploidy, *= diploid), (B) Abundance for each type of TE, and (C) genome size. Red circles indicated those taxa not included in regression analyses.

To test the accuracy of clustering of whole genome shotgun data to assess transposon content, we baselined the Transposome method against several of the published genomes in the Brassicales that were sampled in this study. Our estimates were largely concordant with those in *Carica papaya*, differing by only 1.12% (Nagarajan *et al.* 2008), while they seem to be overestimates compared to the TE abundances found in the *Thlaspi arvense* genome (Dorn *et al.* 2015). The difference in *Thlaspi* estimates is likely the result of the genome assembly and scaffolding strategy, for which the read length and insert size are important for assembling repetitive elements. When insert sizes are small, repetitive elements may collapse, preventing their annotation, evident by the discrepancy between true genome size and assembly size of *T. arvense* (Dorn *et al.* 2015). One benefit of the Transposome and similar RepeatExplorer algorithm approaches is that they do not rely on genome assemblies, for which accurate and complete assembly of repetitive regions is still complicated. Overall, we treated these estimates as a good approximation of TE abundances within their respective genomes.

### Genome size does not correlate to repetitive element content

We next performed regression analysis to test the relationship between genome sizes and their respective TE abundances. The issue with simple regression is that it assumes independence between data points (Felsenstein 1985). To account for phylogenetic relationships between species, and lack of independence thus present in our data, we calculated PIC values for both genome sizes and TE abundances. These values were then used as input for linear regression. Across all the species in this study, there was no significant trend between genome size and repeat content (R^2^=0.0025, p-value=0.698; **Figure 2A**). There was no evidence for a strong overall correlation between genome size and any particular class of transposable elements. However, when we analyze individual families, we identify a strong correlation between genome size and the abundance of *Gypsy* elements in Capparaceae (R^2^=0.99, p-value=0.048; **Figure 2C**). Although, we believe that more extensive sampling is needed to confirm this result, as our sampling only includes four species from the Capparaceae family. The lack of interdependence between genome size and TE abundance was not entirely surprising and has been hinted at by Lysak *et al.* (2009). They found that, while there are exceptions, most Brassicaceae species have relatively small genomes. This was an interesting discovery, as Brassicaceae genomes have undergone multiple polyploidization events and TE proliferation, both of which are expected to lead to an increase in genome size (Jhonston *et al.* 2005; Lysak *et al.* 2009).

**Figure 2.**
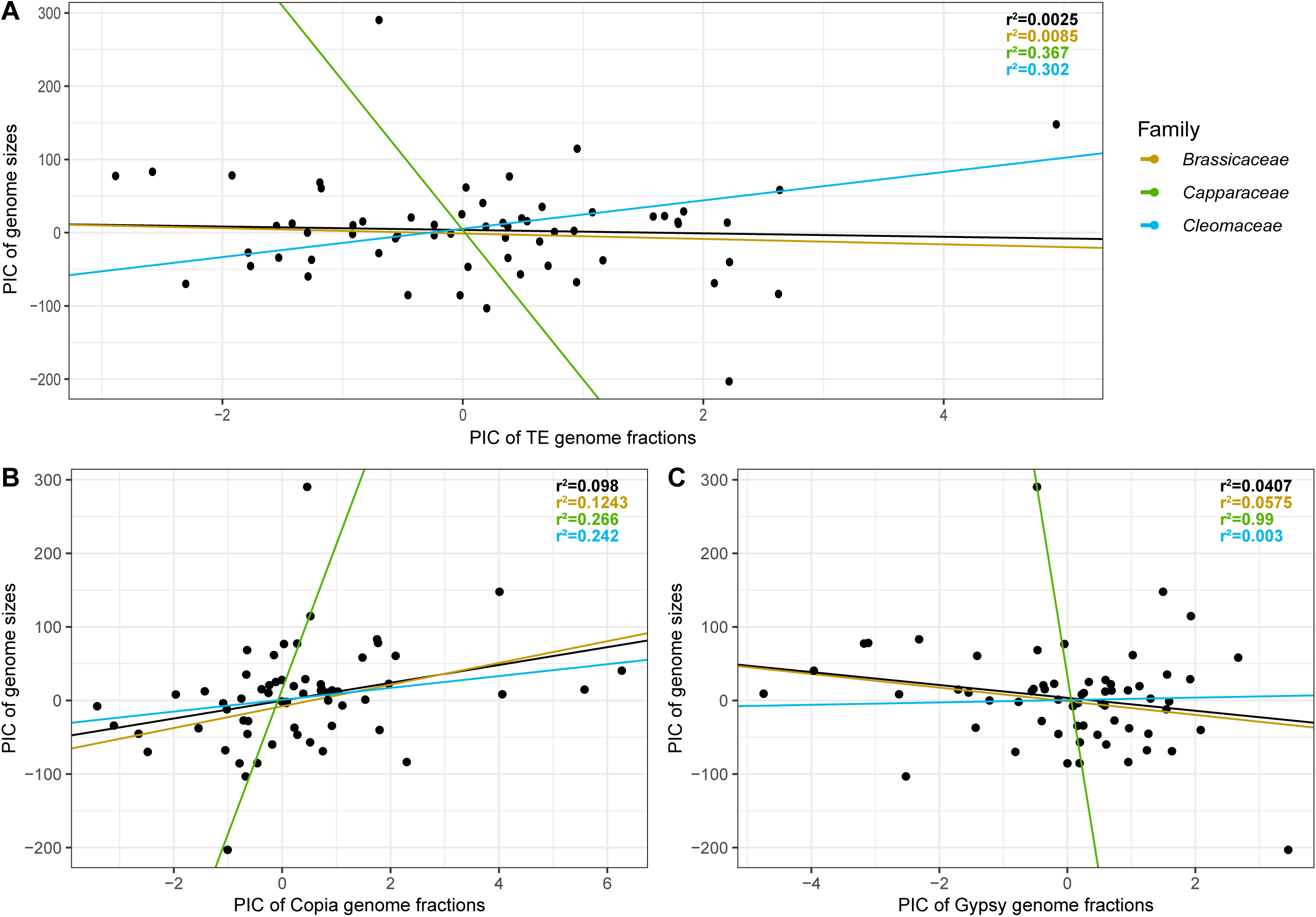
Linear regression analysis of the relationship between genome size and (A) total TE, (B) *Copia* and (C) *Gypsy* element abundance; PIC=phylogenetically independent contrasts.

### TE content does not reflect phylogenetic relationships

To test the congruence of the ASTRAL tree with a repeat clustering based tree, we performed hierarchical clustering of TE abundances for *Copia* and *Gypsy* elements, as well as total TE content. We specifically highlight these superfamilies since previous studies which have used TE abundances to reconstruct phylogenetic trees found that *Copia* and *Gypsy* elements bear the strongest signal (Dodsworth *et al.* 2015; Harkess *et al.* 2016; Dodsworth *et al.* 2017; Vitales *et al.* 2020). However, we were unable to reproduce the species tree using our TE abundance data (**Figure S2**). While a few relationships were established correctly, none of the resulting dendrograms mirrored the tree obtained through ASTRAL using transcriptome data. At the family level, similar to previous publications, we did observe mirroring relationships within some clades (**Figure S3**), while the overall dendrogram was still in conflict with the species phylogeny (Dodsworth *et al.* 2015; Vitales *et al.* 2020). Overall, we observed more agreement within genera, with the level of conflict increasing in higher taxonomic ranks, resulting in a poorly resolved tree across the Brassicales. Our data indicate that this approach does not have enough resolution to elucidate the complex evolutionary history of the Brassicales order.

While some of the disagreement between the species phylogeny and TE abundance analyses could come from the different methods used, we speculate that, on a large scale, other factors driving genome evolution in the Brassicales dilute the phylogenetic signal coming from TE abundances. One possibility is related to certain technical compromises that were necessary in order to run such a large number of species in Transposome. We did not attempt to pool all 71 species into a single clustering analysis, such as performed in smaller species groups like eight species of *Asparagus* (Harkess *et al.* 2016), six *Nicotiana* species (Dodsworth *et al.* 2015; Dodsworth *et al.* 2017; Vitales *et al.* 2020), and nine *Fabeae* species (Dodsworth *et al.* 2015; Vitales *et al.* 2020). Similarly, we did not adjust for genome size due to the large number of sampled species and drastic genome size differences across the samples. Scaling would here result in great underrepresentation of species with smaller genomes in the dataset, diminishing our ability to identify less abundant repeat types. Other studies which have identified phylogenetic signals in their TE content have used much smaller sample sizes and have a much narrower focus on taxa of study (Dodsworth *et al.* 2015; Harkess *et al.* 2016; Dodsworth *et al.* 2017; Vitales *et al.* 2020). Further, as described in the documentation of RepeatExplorer (http://repeatexplorer.org/?page_id=179), our sample size should be able to well-represent highly abundant TEs, and genome size adjustments have more significance for detection of low-copy repeats.

### Polyploidy is not correlated to shifts in TE abundance

Previous work, restricted to neopolyploids, has indicated that WGD and TE abundance, both expansion and loss, are correlated (Ågren *et al.* 2016; Parisod *et al.* 2010; Vicient and Casacuberta, 2017). Here, we did not recover any evidence to support such a correlation across the Brassicales, a much older timescale (**Figure 3**). For total TE abundance, we identified a shift toward a higher abundance in a clade of the Cleomaceae comprising the genus *Polanisia* and three species of *Cleome.* Surprisingly, this clade is sister to the clade recently characterized by the Cleomaceae specific WGD, Th – α (Mabry *et al.* 2020). For *Gypsy* elements alone, we identified a single shift in *Lunaria annua;* this was unsurprising, as in **Figure 1**, clear differences in the proportion of these elements can be seen. Two shifts were identified for *Copia* abundance, one for *Hesperis matronalis*, which has a very large genome, and one for the *Polanisia* clade. In the *Polanisia* clade, *Copia* elements comprised, on average, 31.5% of the genome, which can be compared to its sister clade, in which *Copia* elements make up on average 8.5% of the genome. For all three TE categories, we did not recover any shifts that overlap with known polyploidy events in the Brassicales (**Figure 3**).

**Figure 3.**
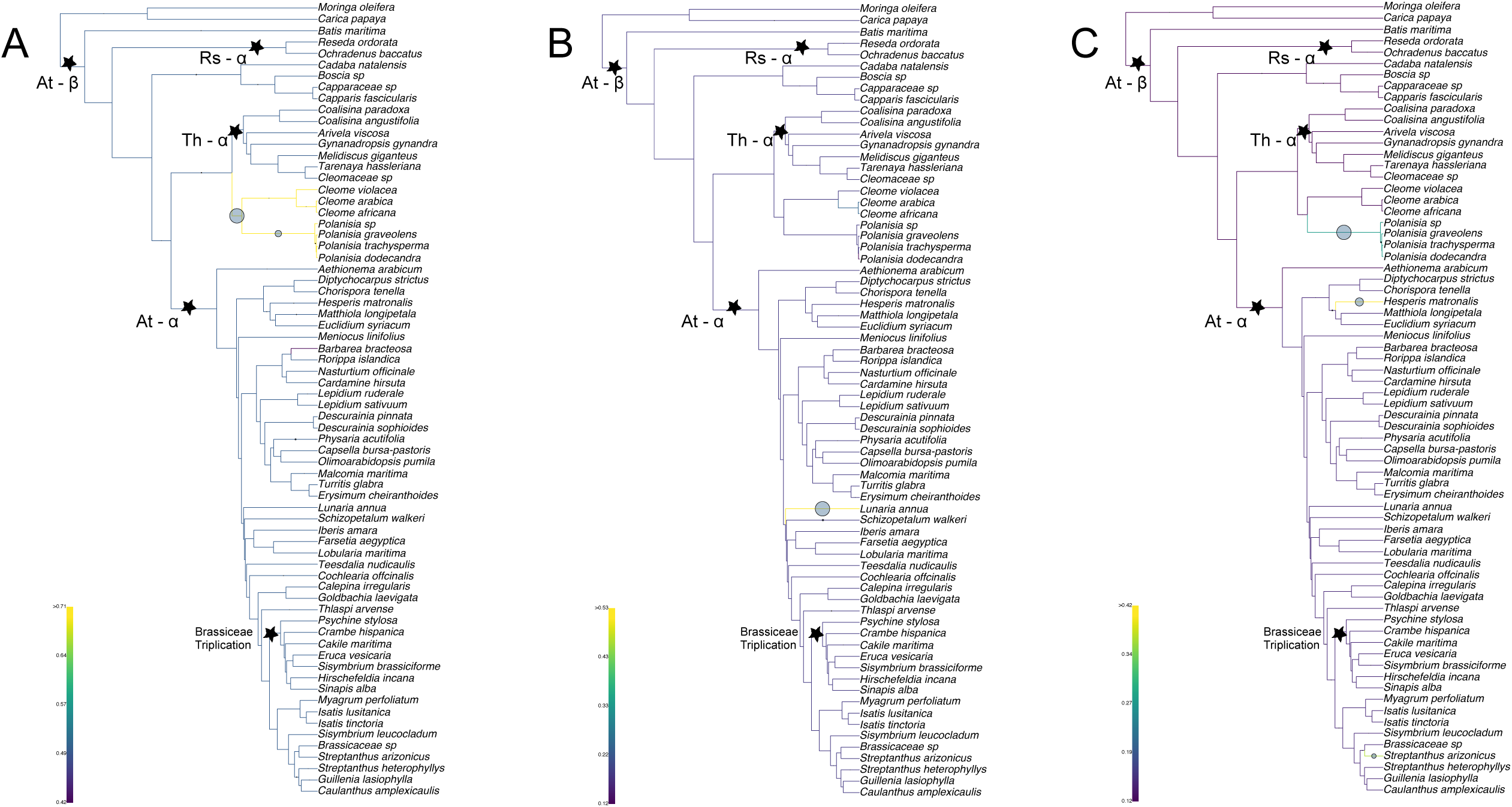
Bayou analysis for detecting shifts in transposable element abundance. (A) Heatmap of total TE, (B) *Gyspy*, and (C) *Copia* abundance with identified shifts plotted. Only shifts with a posterior probability of 0.3 are plotted. Circle size corresponds to the posterior probability of having a shift. Whole genome duplications are denoted by black stars with named events indicated.

To further test the hypothesis that WGD and TE abundances are correlated, we constrained our analyses to implicitly test for shifts at known WGD events using OUwie. All analyses indicated that there is no correlation between TE proliferation and WGD (**Table S4**). Specifically, when testing for correlations between At – α at the base of the Brassicaceae versus all Brassicales for total TE abundance, the BMS model (Brownian motion, with different rate parameters for each state on a tree) was assigned the most weight compared to the other models tested. The BMS model suggested that there is a single optima that the taxa are moving toward, albeit with different rates (Sigma^2 = 0.0001378363 for no WGD and 0.0003516384 with the WGD). The OU 1 model (Ornstein-Uhlenbeck model with a single optimum for all species) was weighted highest for both *Gyspy* and *Copia* elements when constraining the selection regium at At – α, again suggesting that there is a single optima that the plants are moving toward, regardless if they have experienced the At – α event or not. For the tribe Brassiceae WGT, the OU 1 event was again weighted highest when compared to the other models, meaning that for this event, taxa with or without the WGT are moving toward a single optima in TE abundance (total TE, *Gyspy*, and *Copia*). For the Th – α event of the Cleomaceae, the same conclusion was drawn, with total TE abundance recovering the BM1 model with the most weight and *Gyspy* and *Copia* analyses weighing the OU1 model highest, taxa with or without the WGD were again recovered as moving toward the same optima. This was somewhat surprising, as Bayou did recover a shift for one clade of Cleomaceae when testing total TEs, and then a shift was observed for just the *Polanisa* clade for *Copia* abundance alone. Since previous studies that have shown a correlation between WGD and TE abundance have typically used neopolyploid species, we further subsetted our dataset to include only taxa for which we had recently sequenced the genome (*unplublished data*) and therefore could confidently identify ploidy, allowing us to test neopolyploids vs known diploids. These analyses also found no correlation between TE abundance and WGD (**Table S4**). The BM1 model was most highly weighted for Total TE and *Gyspy* while the OU1 model was most highly rated for *Copia*.

Although it has been hypothesized that WGD may be correlated with genome size (Ågren *et al.* 2016; Vicient and Casacuberta 2017), we did not recover support for this correlation within the Brassicales, a group characterized by multiple WGD events. One hypothesis for these patterns is that the genomes that share the three events tested here (At – α, Th – α, and the tribe Brassiceae WGT) have already diploidized. However, we were especially surprised to find no correlation between TE and WGD when testing neopolyploidy events using the reduced subset of data. Many of the studies which have found evidence for this correlation have typically assessed a single species with a recent WGD, for example Petit *et al.* (2010) found TE proliferation after polyploidization in tobacco, Madlung *et al.* (2005) show increased activity of several transposons in newly formed allopolyploid *Arabidopsis,* as did Kashkush *et al.* (2003) and Lopes *et al.* (2013) in wheat and coffee respectively, with many more additional examples of polyploidy correlated with TEs composed by Vicient and Casacuberta (2017). Yet, Ågren *et al.* (2016) suggest that for *Capsella bursa-pastoris,* TE abundance increased due to relaxed selection, while Hu *et al.* (2010) and Charles *et al.* (2008) found no evidence of proliferation of TEs after WGD in cotton and wheat, respectively. Looking more broadly for support for this WGD – TE abundance correlation, Staton and Burke (2015) assessed 15 taxa across the Asteraceae. Although they place WGD events on their phylogeny, they note that further work is needed to test if this is a true correlation. It seems that although there is still this predominant theory that WGD and TE abundance are correlated, researchers have begun to appreciate that the story is much more complex (Parisod *et al.* 2010; Sarilar *et al.* 2013).

In the absence of a technical explanation, another possibility relates to the genome stasis of species in the Brassicales following polyploidy and subsequent diploidization. That is, there could exist a mechanism(s) to suppress repetitive element proliferation and diversification enabling the tape of evolution to be “replayed” (Bird *et al.* 2019). For example, when testing resynthesized polyploid *Brassica napus* lines, Bird *et al.* (2019), found that the same parental subgenome was consistently more dominantly expressed in all lines and generations. The subgenomes of wheat were also found to be surprisingly static after hybridization (Wicker *et al.* 2018). Overall, results here and from others cited within provide support for a type of punctuated equilibria, where evolutionary development is marked by isolated episodes of rapid change as noted in many crop polyploid species, between long periods of little or no change, as a way to explain the patterns we see here (Zeh, Zeh, and Ishida 2009).

Using 71 taxa across the Brassicales, we found no evidence for either a phylogenetic signal from TEs or a correlation between WGD and TE abundance. This study is the first to assess TE abundance across an entire plant order with this many samples. We suggest that although TE abundance may follow phylogenetic signals at shallow phylogenetic levels, it should be used with caution for determining relationships at deeper nodes of a phylogeny. We also suggest that, although TE abundance may be driven by WGD at short time scales, TE expansion does not leave an overall lasting imprint on a genome and that TE purging mechanisms, such as intrastrand homologous recombination and illegitimate recombination, work efficiently to bring genomes to stability (Hawkins *et al.* 2009). As the cost of genomes continues to decrease, the opportunities to test these patterns by annotating TEs in multiple assembled genomes per family will be possible, although hinging on computational limitations. These analyses paired with others which test for patterns of TE evolution in other groups of organisms, such as other plant orders and even larger groups across the tree of life will hopefully provide insight to further understand the C-value paradox.

## ACKNOWLEDGEMENTS

The authors would like to thank Predreg Lazic from the research computing support services at the University of Missouri, Josh Rothhaupt from the Data Science team at the Donald Danforth Plant Science Center, Dr. Paul Blischak from the University of Arizona, and Christian Concepcion for their assistance in both getting software to run and help in understanding models, and code development. We thank our Molecular and Network Evolution course at the University of Missouri, which enabled our project and introduced the co-first authors to one another. We thank both our funding and computational resources; Department of Energy Defense Threat Reduction Agency (HDTRA 1-16-1-0048), National Science Foundation (IOS 1339156), and the Research Computing Support Services (RCSS) at the University of Missouri. Lastly, we thank our anonymous reviews for their comments and edits of our manuscript.

## AUTHOR CONTRIBUTIONS

AB, MEM, AEH, PPE, MES, GCC, BCM, and JCP designed the study. MEM grew, sampled, and collected RNA/DNA from plants. AB conducted TE annotation, regression analyses and phylogenetic comparisons of TE-derived phylogeny to species tree. MEM performed species tree inference and comparative genomic analyses. AEH and GCC helped with analyses. AB and MEM wrote the manuscript. All authors provided feedback and helped shape the final manuscript.

## SUPPLEMENTAL MATERIAL

**Supplemental Figure 1.** Proportion plots of transposable element abundance scaled to 100%. (A) phylogeny of taxa, (B) by superfamily, (C) by order.

**Supplemental Figure 2.** Comparison of phylogenetic relationships across Brassicales in the ASTRAL tree obtained from transcriptome data (left) and based on hierarchical clustering of TE abundances (right).

**Supplemental Figure 3.** Comparison of phylogenetic relationships in the ASTRAL tree obtained from transcriptome data (black) and based on hierarchical clustering of TE abundances (red) in (A) Brassicaceae, (B) Cleomaceae, and (C) Capparaceae.

**Supplemental Table 1.** Taxon sampling with genome size, chromosome counts, and transposable element abundance. Chromosome counts as in http://legacy.tropicos.org/Project/IPCN. Asterisks (*) indicate samples with known ploidy level.

**Supplemental Table 2.** List of mitochondrial and chloroplast sequences used for data filtering.

**Supplemental Table 3.** Data processing summary.

**Supplemental Table 4.** OUwie weighted AICc scores. Tests conducted are specified below as selection regime ~ modelled phenotype. BM1; single-rate Brownian motion, BMS; Brownian motion with different rate parameters for each state on a tree, OU1; Ornstein-Uhlenbeck model with a single optimum for all species, OUM; Ornstein-Uhlenbeck model with different state means and a single alpha and sigma^2 acting all selective regimes, OUMV; Ornstein-Uhlenbeck model that assumes different state means as well as multiple sigma^2, OUMA; Ornstein-Uhlenbeck model that assumes different state means as well as multiple alpha, OUMVA; Ornstein-Uhlenbeck model that assumes different state means as well as multiple alpha and sigma^2 per selective regime. Asterisk (*) indicated analyses which returned the warning, “You might not have enough data to fit this model well”.

