## Supplemental Figures and Tables for "Surprising amount of stasis in repetitive genome content across the Brassicales"

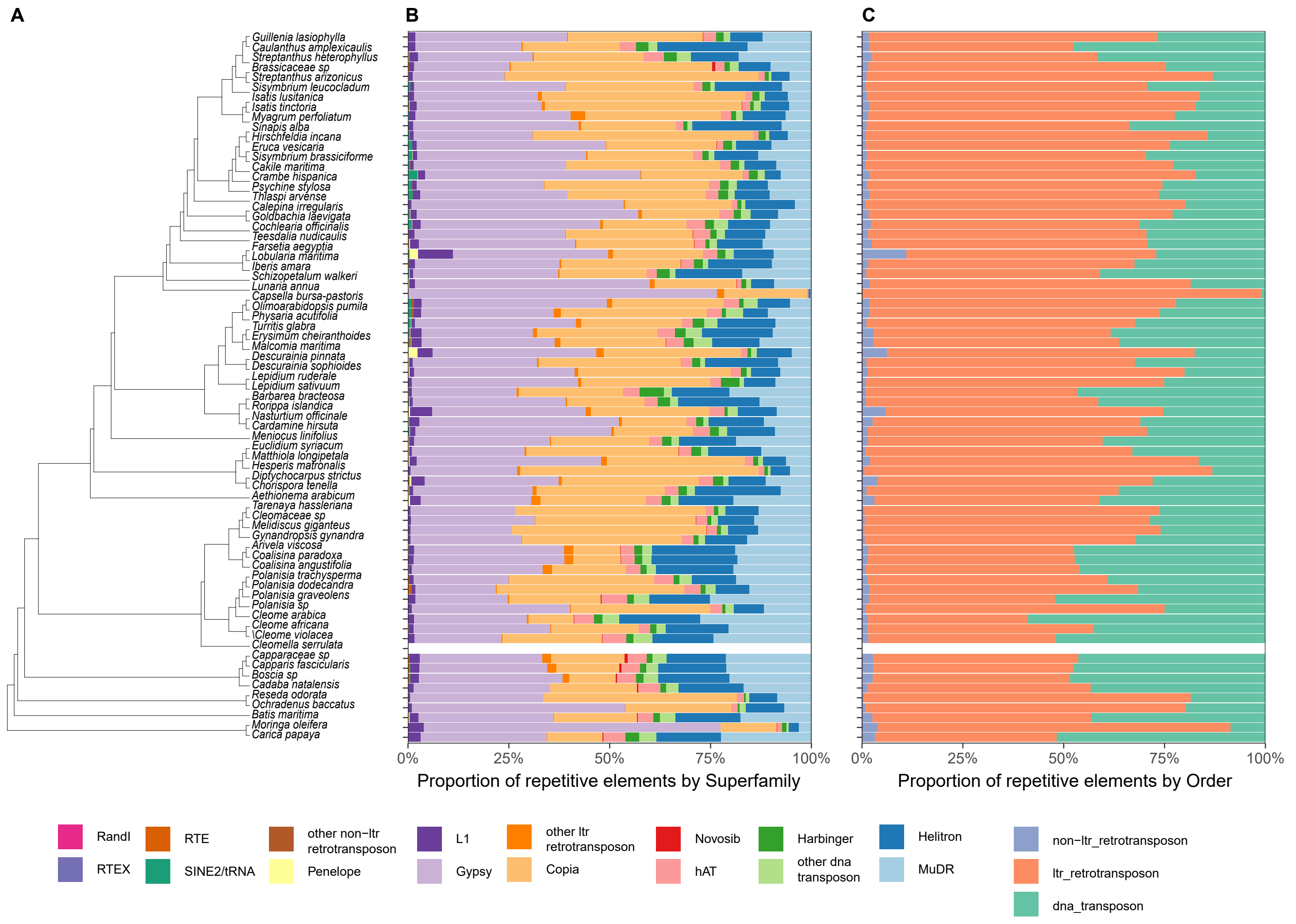


**Supplemental Figure 1.** Proportion plots of transposable element abundance scaled to 100%. (A) phylogeny of taxa, (B) by superfamily, (C) by order.

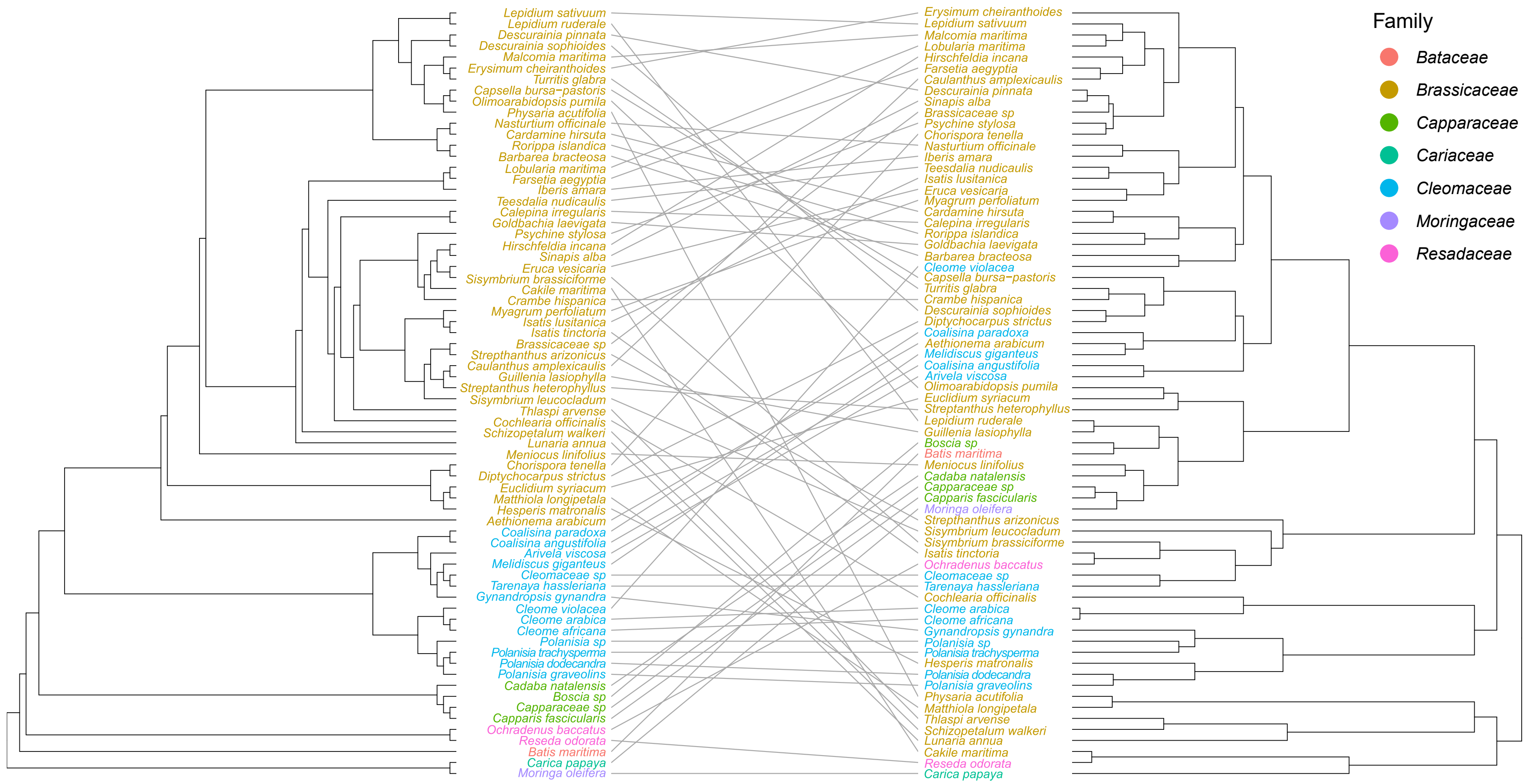


**Supplemental Figure 2.** Comparison of phylogenetic relationships across Brassicales in the ASTRAL tree obtained from transcriptome data (left) and based on hierarchical clustering of TE abundances (right).

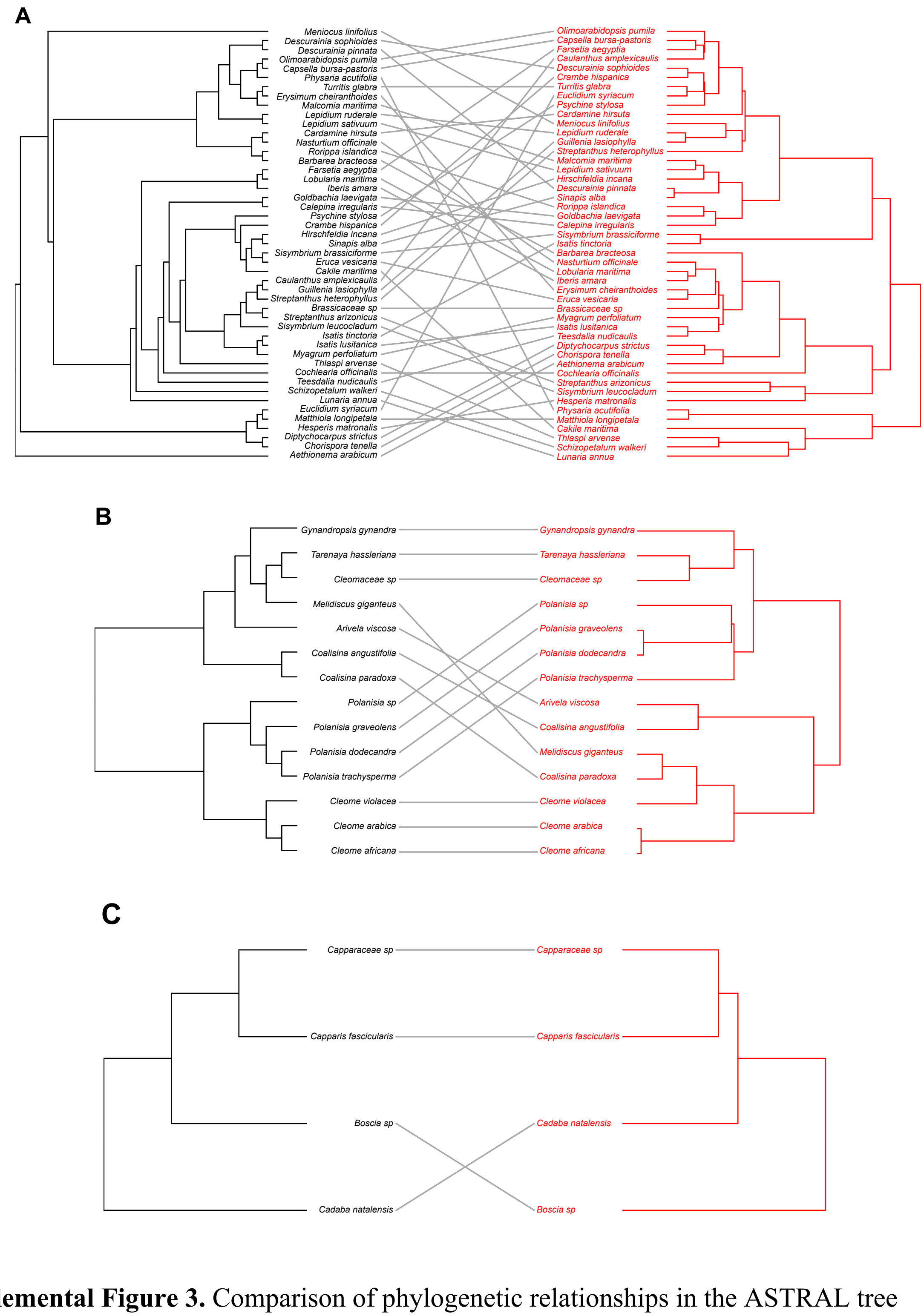


**Supplemental Figure 3.** Comparison of phylogenetic relationships in the ASTRAL tree obtained from transcriptome data (black) and based on hierarchical clustering of TE abundances (red) in (A) Brassicaceae, (B) Cleomaceae, and (C) Capparaceae.

**Supplemental Table 1.** Taxon sampling with genome size, chromosome counts, and repetitive element abundance. Chromosome counts as in http://legacy.tropicos.org/Project/IPCN. Asterisks (*) indicate samples with known ploidy level.

| Family | Species | Genome Size (Mbp) | Chromosome count (n) | Copia | Gypsy | Harbinger | hAT | Helitron | L1 | MuDR | other DNA transposon | other ltr retrotransposon | other non-ltr retrotransposon | Penelope | SINE2/tRNA | Novosib | RTE | RTEX | RandI | Total_RE |
|---|---|---|---|---|---|---|---|---|---|---|---|---|---|---|---|---|---|---|---|---|
| Bataceae | *Batis maritima* | 356.97 | 11 | 0.0734 | 0.1192 | 0.0058 | 0.0138 | 0.0571 | 0.0075 | 0.0625 | 1.38E-02 | 4.17E-04 | 0.00E+00 | 8.34E-04 | 0.00E+00 | 4.17E-04 | 8.34E-04 | 0.00E+00 | 0.00E+00 | 0.3556 |
| Brassicaceae | *Aethionema arabicum** | 933.99 | 11 | 0.1557 | 0.1632 | 0.0129 | 0.0235 | 0.0811 | 0.0163 | 0.115 | 1.23E-02 | 1.36E-02 | 4.04E-04 | 2.16E-03 | 2.70E-04 | 0.00E+00 | 0.00E+00 | 0.00E+00 | 0.00E+00 | 0.5964 |
| Brassicaceae | *Meniocus linifolius* | 899.76 | | 0.0997 | 0.1382 | 0.0098 | 0.0132 | 0.0581 | 0.0049 | 0.0768 | 7.48E-03 | 1.23E-03 | 3.79E-04 | 5.68E-04 | 0.00E+00 | 9.47E-05 | 2.84E-04 | 9.47E-05 | 0.00E+00 | 0.4107 |
| Brassicaceae | *Barbarea bracteosa* | 268.95 | 8 | 0.1068 | 0.2629 | 0.0124 | 0.0218 | 0.0637 | 0.0073 | 0.0491 | 1.13E-02 | 3.59E-03 | 7.98E-05 | 5.59E-04 | 1.92E-03 | 0.00E+00 | 3.19E-04 | 0.00E+00 | 0.00E+00 | 0.5417 |
| Brassicaceae | *Cakile maritima** | 1383.87 | 9 | 0.1463 | 0.3109 | 0.0133 | 0.0082 | 0.0225 | 0.0101 | 0.0444 | 1.03E-02 | 1.33E-03 | 0.00E+00 | 6.66E-04 | 1.38E-02 | 0.00E+00 | 0.00E+00 | 0.00E+00 | 0.00E+00 | 0.5817 |
| Brassicaceae | *Calepina irregularis* | 254.28 | 14 | 0.0957 | 0.2056 | 0.0104 | 0.0213 | 0.0483 | 0.0095 | 0.0471 | 1.57E-02 | 2.97E-03 | 6.20E-04 | 2.48E-04 | 3.72E-03 | 0.00E+00 | 2.48E-04 | 1.24E-04 | 0.00E+00 | 0.4617 |
| Brassicaceae | *Capsella bursa-pastoris* | 420.54 | 8,16 | 0.1275 | 0.1617 | 0.0116 | 0.0218 | 0.0571 | 0.0122 | 0.0631 | 2.27E-02 | 7.04E-03 | 8.80E-04 | 6.40E-04 | 1.84E-03 | 3.20E-04 | 8.00E-04 | 1.60E-04 | 0.00E+00 | 0.4893 |
| Brassicaceae | *Cardamine hirsuta* | 489 | 7,8 | 0.0911 | 0.1797 | 0.0145 | 0.0153 | 0.0953 | 0.0036 | 0.0608 | 9.44E-03 | 1.42E-03 | 3.78E-04 | 2.83E-04 | 4.72E-04 | 0.00E+00 | 7.55E-04 | 0.00E+00 | 0.00E+00 | 0.4731 |
| Brassicaceae | *Caulanthus amplexicaulis** | 484.11 | | 0.1521 | 0.1708 | 0.0084 | 0.0146 | 0.0362 | 0.0074 | 0.0549 | 7.60E-03 | 1.36E-03 | 3.90E-04 | 0.00E+00 | 0.00E+00 | 9.74E-05 | 5.84E-04 | 0.00E+00 | 0.00E+00 | 0.4544 |
| Brassicaceae | *Chorispora tenella* | 366.75 | 7 | 0.174 | 0.1702 | 0.0126 | 0.0175 | 0.0471 | 0.0167 | 0.0578 | 7.26E-03 | 3.98E-03 | 6.96E-04 | 2.59E-03 | 8.95E-04 | 9.94E-05 | 3.98E-04 | 0.00E+00 | 0.00E+00 | 0.5118 |
| Brassicaceae | *Cochlearia offcinalis* | 734 | 12 | 0.1941 | 0.231 | 0.009 | 0.0271 | 0.0615 | 0.0082 | 0.0707 | 1.32E-02 | 1.12E-03 | 3.75E-04 | 2.50E-04 | 0.00E+00 | 5.00E-04 | 9.99E-04 | 0.00E+00 | 0.00E+00 | 0.6181 |
| Brassicaceae | *Crambe hispanica** | 660.15 | 15,30 | 0.1942 | 0.1496 | 0.0099 | 0.0126 | 0.0362 | 0.0053 | 0.0513 | 9.91E-03 | 5.08E-04 | 2.54E-04 | 2.54E-04 | 4.49E-03 | 0.00E+00 | 1.69E-04 | 8.47E-05 | 0.00E+00 | 0.4749 |
| Brassicaceae | *Descurainia pinnata** | 371.64 | 7 | 0.1866 | 0.1963 | 0.0069 | 0.0124 | 0.0356 | 0.0056 | 0.0383 | 5.44E-03 | 4.57E-03 | 3.89E-04 | 1.36E-03 | 9.72E-05 | 9.72E-05 | 2.91E-04 | 0.00E+00 | 0.00E+00 | 0.4939 |
| Brassicaceae | *Descurainia sophioides** | 195.6 | 7,14 | 0.175 | 0.1532 | 0.0096 | 0.014 | 0.0897 | 0.0045 | 0.0412 | 6.16E-03 | 2.22E-03 | 1.01E-04 | 1.01E-03 | 3.03E-04 | 0.00E+00 | 1.01E-04 | 0.00E+00 | 0.00E+00 | 0.4973 |
| Brassicaceae | *Diptychocarpus strictus** | 557.46 | 7 | 0.1625 | 0.1509 | 0.0106 | 0.0168 | 0.1085 | 0.0051 | 0.0385 | 1.05E-02 | 4.98E-03 | 4.24E-04 | 4.24E-04 | 3.18E-04 | 1.06E-04 | 3.18E-04 | 0.00E+00 | 0.00E+00 | 0.5101 |
| Brassicaceae | *Eruca vesicaria** | 709.05 | 11 | 0.1464 | 0.2511 | 0.012 | 0.008 | 0.0465 | 0.0053 | 0.0529 | 5.58E-03 | 4.04E-04 | 2.42E-04 | 0.00E+00 | 5.33E-03 | 1.62E-04 | 3.23E-04 | 1.62E-04 | 0.00E+00 | 0.5345 |
| Brassicaceae | *Erysimum cheiranthoides* | 420.54 | 8 | 0.1424 | 0.2372 | 0.0056 | 0.0196 | 0.041 | 0.0093 | 0.0274 | 1.84E-02 | 7.27E-03 | 2.32E-03 | 1.83E-04 | 5.01E-03 | 2.44E-04 | 3.05E-04 | 0.00E+00 | 0.00E+00 | 0.5164 |
| Brassicaceae | *Euclidium syriacum** | 298.29 | 7 | 0.1686 | 0.1255 | 0.0105 | 0.014 | 0.0591 | 0.0027 | 0.0554 | 7.99E-03 | 2.00E-03 | 5.26E-04 | 4.20E-04 | 1.05E-04 | 4.20E-04 | 4.20E-04 | 0.00E+00 | 0.00E+00 | 0.4477 |
| Brassicaceae | *Farsetia aegyptica* | 2004.9 | 24 | 0.142 | 0.1715 | 0.0101 | 0.0155 | 0.0753 | 0.0067 | 0.0471 | 6.58E-03 | 2.55E-03 | 8.22E-05 | 2.47E-04 | 8.22E-04 | 8.22E-05 | 3.29E-04 | 8.22E-05 | 0.00E+00 | 0.479 |
| Brassicaceae | *Goldbachia laevigata* | 547.68 | 14 | 0.0885 | 0.2542 | 0.0076 | 0.0168 | 0.0314 | 0.0072 | 0.0382 | 1.14E-02 | 4.31E-03 | 1.96E-04 | 1.96E-04 | 2.06E-03 | 2.94E-04 | 4.89E-04 | 0.00E+00 | 0.00E+00 | 0.463 |
| Brassicaceae | *Guillenia lasiophylla* | 621.03 | | 0.0908 | 0.0997 | 0.0116 | 0.0153 | 0.0846 | 0.0066 | 0.06 | 8.36E-03 | 1.38E-03 | 2.30E-04 | 1.53E-04 | 0.00E+00 | 1.53E-04 | 0.00E+00 | 0.00E+00 | 0.00E+00 | 0.3788 |
| Brassicaceae | *Hesperis matronalis* | 3261.63 | 7 | 0.395 | 0.1763 | 0.0056 | 0.0097 | 0.0321 | 0.0038 | 0.0354 | 4.32E-03 | 4.75E-03 | 0.00E+00 | 0.00E+00 | 7.38E-04 | 0.00E+00 | 0.00E+00 | 0.00E+00 | 0.00E+00 | 0.6677 |
| Brassicaceae | *Hirschefeldia incana* | 552.57 | 7 | 0.1222 | 0.1947 | 0.0072 | 0.0107 | 0.0505 | 0.0056 | 0.0616 | 6.36E-03 | 1.47E-03 | 4.71E-04 | 5.89E-05 | 4.59E-03 | 0.00E+00 | 1.18E-04 | 5.89E-05 | 0.00E+00 | 0.4655 |
| Brassicaceae | *Iberis amara** | 831.3 | 7 | 0.1167 | 0.1951 | 0.0173 | 0.014 | 0.0897 | 0.0046 | 0.0929 | 7.37E-03 | 2.05E-03 | 2.12E-04 | 4.96E-04 | 1.35E-03 | 7.08E-05 | 2.12E-04 | 0.00E+00 | 0.00E+00 | 0.542 |
| Brassicaceae | *Isatis lusitanica* | 381.42 | 14 | 0.1909 | 0.2181 | 0.0067 | 0.0139 | 0.0602 | 0.0085 | 0.0359 | 9.47E-03 | 1.99E-02 | 1.39E-04 | 0.00E+00 | 1.39E-03 | 0.00E+00 | 2.79E-04 | 0.00E+00 | 0.00E+00 | 0.5654 |
| Brassicaceae | *Isatis tinctoria** | 616.14 | 8,16 | 0.2403 | 0.1527 | 0.0044 | 0.0102 | 0.0346 | 0.0086 | 0.0271 | 8.30E-03 | 4.12E-03 | 4.31E-04 | 5.54E-04 | 9.84E-04 | 6.15E-05 | 1.85E-04 | 0.00E+00 | 0.00E+00 | 0.4925 |

| | | | | | | | | | | | | | | | | | | | | |
|---|---|---|---|---|---|---|---|---|---|---|---|---|---|---|---|---|---|---|---|---|
| Brassicaceae | *Lepidium ruderale* | 371.64 | 12,16 | 0.1015 | 0.1015 | 0.0234 | 0.0158 | 0.0561 | 0.003 | 0.0795 | 7.69E-03 | 2.53E-03 | 0.00E+00 | 3.89E-04 | 0.00E+00 | 0.00E+00 | 3.89E-04 | 9.73E-05 | 0.00E+00 | 0.3918 |
| Brassicaceae | *Lepidium sativuum** | 591.69 | 12 | 0.1574 | 0.2023 | 0.0225 | 0.0131 | 0.038 | 0.0044 | 0.0439 | 5.46E-03 | 3.64E-03 | 1.07E-04 | 1.07E-04 | 0.00E+00 | 0.00E+00 | 4.28E-04 | 0.00E+00 | 0.00E+00 | 0.4912 |
| Brassicaceae | *Lobularia maritima* | 312.96 | 15 | 0.1145 | 0.1981 | 0.01 | 0.0185 | 0.0507 | 0.0447 | 0.0483 | 1.12E-02 | 6.33E-03 | 0.00E+00 | 1.09E-02 | 1.45E-03 | 2.64E-04 | 7.92E-04 | 0.00E+00 | 0.00E+00 | 0.5158 |
| Brassicaceae | *Lunaria annua** | 498.78 | 8 | 0.1296 | 0.4807 | 0 | 0.002 | 0.0024 | 0 | 0.0016 | 0.00E+00 | 1.05E-02 | 0.00E+00 | 0.00E+00 | 0.00E+00 | 0.00E+00 | 0.00E+00 | 0.00E+00 | 0.00E+00 | 0.6269 |
| Brassicaceae | *Malcomia maritima** | 312.96 | 7 | 0.1218 | 0.1949 | 0.0136 | 0.0129 | 0.07 | 0.0048 | 0.0436 | 1.62E-02 | 6.46E-03 | 2.75E-04 | 1.38E-04 | 3.16E-03 | 0.00E+00 | 2.75E-04 | 0.00E+00 | 0.00E+00 | 0.4882 |
| Brassicaceae | *Matthiola longipetala* | 1599.03 | 7 | 0.2362 | 0.316 | 0.0086 | 0.0146 | 0.0395 | 0.0118 | 0.0432 | 7.71E-03 | 9.56E-03 | 7.14E-04 | 1.28E-03 | 0.00E+00 | 0.00E+00 | 7.14E-04 | 0.00E+00 | 0.00E+00 | 0.69 |
| Brassicaceae | *Myagrum perfoliatum** | 352.08 | 16 | 0.1296 | 0.2265 | 0.0053 | 0.01 | 0.1219 | 0.006 | 0.041 | 6.75E-03 | 3.45E-03 | 1.57E-04 | 0.00E+00 | 7.85E-04 | 1.57E-04 | 3.14E-04 | 0.00E+00 | 0.00E+00 | 0.552 |
| Brassicaceae | *Nasturtium officinale* | 430.32 | 15 | 0.1584 | 0.206 | 0.0046 | 0.0198 | 0.052 | 0.029 | 0.0463 | 1.35E-02 | 6.96E-03 | 1.31E-04 | 2.23E-03 | 3.94E-04 | 2.63E-04 | 2.63E-04 | 1.31E-04 | 0.00E+00 | 0.54 |
| Brassicaceae | *Olimoarabidopsis pumila* | 410.76 | 5 | 0.132 | 0.1227 | 0.0119 | 0.0191 | 0.0773 | 0.012 | 0.0425 | 1.81E-02 | 4.07E-03 | 9.20E-04 | 3.94E-04 | 8.54E-04 | 6.57E-05 | 5.26E-04 | 6.57E-05 | 0.00E+00 | 0.4424 |
| Brassicaceae | *Physaria acutifolia* | 2322.75 | | 0.2462 | 0.2936 | 0.0066 | 0.0115 | 0.0627 | 0.0274 | 0.0348 | 1.04E-02 | 1.27E-02 | 6.49E-05 | 1.62E-02 | 0.00E+00 | 0.00E+00 | 6.49E-04 | 1.30E-04 | 0.00E+00 | 0.723 |
| Brassicaceae | *Sisymbrium leucocladum* | 498.78 | 8 | 0.2872 | 0.1753 | 0.0099 | 0.0098 | 0.0325 | 0.0072 | 0.0331 | 7.51E-03 | 5.03E-03 | 2.40E-04 | 0.00E+00 | 8.79E-04 | 3.19E-04 | 2.40E-04 | 0.00E+00 | 0.00E+00 | 0.5693 |
| Brassicaceae | *Psychine stylosa* | 518.34 | 16 | 0.1695 | 0.1805 | 0.0121 | 0.0147 | 0.0432 | 0.0085 | 0.0509 | 7.72E-03 | 2.86E-04 | 4.77E-04 | 9.53E-05 | 5.62E-03 | 0.00E+00 | 2.86E-04 | 0.00E+00 | 0.00E+00 | 0.4938 |
| Brassicaceae | *Rorippa islandica** | 567.24 | 10 | 0.0786 | 0.2426 | 0.0082 | 0.0121 | 0.0672 | 0.0123 | 0.0574 | 6.64E-03 | 3.50E-03 | 1.21E-04 | 3.62E-04 | 6.64E-04 | 0.00E+00 | 1.81E-04 | 0.00E+00 | 0.00E+00 | 0.4899 |
| Brassicaceae | *Schizopetalum walkeri* | 1775.07 | 12 | 0.1289 | 0.3687 | 0.0073 | 0.0074 | 0.0363 | 0.009 | 0.0584 | 7.69E-03 | 7.84E-03 | 0.00E+00 | 2.22E-03 | 0.00E+00 | 1.48E-04 | 1.48E-04 | 0.00E+00 | 0.00E+00 | 0.634 |
| Brassicaceae | *Sinapis alba** | 557.46 | 7 | 0.1894 | 0.1884 | 0.0103 | 0.0137 | 0.0391 | 0.0038 | 0.0434 | 6.77E-03 | 4.90E-04 | 3.92E-04 | 1.96E-04 | 2.45E-03 | 9.81E-05 | 1.96E-04 | 9.81E-05 | 0.00E+00 | 0.4989 |
| Brassicaceae | *Sisymbrium brassiciforme* | NA | 14 | 0.2708 | 0.1461 | 0.0091 | 0.0063 | 0.0223 | 0.004 | 0.0292 | 4.39E-03 | 4.18E-04 | 1.04E-04 | 5.22E-05 | 2.40E-03 | 0.00E+00 | 3.13E-04 | 5.22E-05 | 0.00E+00 | 0.4955 |
| Brassicaceae | *Brassicaceae sp** | 1036.68 | 14 | 0.1639 | 0.1945 | 0.0108 | 0.0112 | 0.0858 | 0.0041 | 0.0377 | 5.69E-03 | 4.90E-04 | 2.94E-04 | 0.00E+00 | 2.65E-03 | 9.81E-05 | 2.94E-04 | 1.96E-04 | 0.00E+00 | 0.5176 |
| Brassicaceae | *Streptanthus arizonicus* | 1090.47 | | 0.3666 | 0.1325 | 0.0053 | 0.0089 | 0.0259 | 0.0059 | 0.0318 | 3.97E-03 | 5.68E-04 | 2.27E-04 | 7.95E-04 | 0.00E+00 | 0.00E+00 | 2.27E-04 | 0.00E+00 | 0.00E+00 | 0.5826 |
| Brassicaceae | *Streptanthus heterophyllys* | 748.17 | 18 | 0.1889 | 0.0898 | 0.0052 | 0.009 | 0.0298 | 0.0046 | 0.0384 | 7.93E-03 | 1.00E-03 | 7.70E-05 | 7.70E-05 | 0.00E+00 | 2.77E-03 | 1.00E-03 | 0.00E+00 | 7.70E-05 | 0.3787 |
| Brassicaceae | *Teesdalia nudicaulis* | 557.46 | 7 | 0.1623 | 0.2155 | 0.0056 | 0.0154 | 0.0625 | 0.0117 | 0.0674 | 1.02E-02 | 1.70E-03 | 9.26E-04 | 1.23E-03 | 0.00E+00 | 1.54E-04 | 9.26E-04 | 0.00E+00 | 0.00E+00 | 0.5555 |
| Brassicaceae | *Thlaspi arvense** | 205.38 | 6 | 0.173 | 0.3441 | 0.0067 | 0.009 | 0.0807 | 0.005 | 0.0264 | 6.04E-03 | 1.56E-03 | 2.08E-04 | 0.00E+00 | 0.00E+00 | 1.04E-04 | 3.12E-04 | 0.00E+00 | 0.00E+00 | 0.653 |
| Brassicaceae | *Turritis glabra* | 919.32 | | 0.1671 | 0.1524 | 0.0046 | 0.0171 | 0.0281 | 0.0084 | 0.0498 | 2.00E-02 | 7.78E-03 | 1.60E-03 | 1.50E-04 | 4.59E-03 | 2.49E-04 | 1.50E-04 | 0.00E+00 | 0.00E+00 | 0.462 |
| Capparaceae | *Boscia sp* | 410.76 | | 0.0452 | 0.1392 | 0.0076 | 0.0183 | 0.0691 | 0.0079 | 0.0791 | 1.37E-02 | 6.31E-03 | 0.00E+00 | 1.77E-04 | 5.89E-05 | 1.18E-03 | 2.36E-03 | 0.00E+00 | 0.00E+00 | 0.3903 |
| Capparaceae | *Cadaba natalensis* | 259.17 | | 0.0935 | 0.1473 | 0.0061 | 0.0239 | 0.0699 | 0.0057 | 0.0735 | 1.39E-02 | 4.74E-04 | 0.00E+00 | 0.00E+00 | 1.18E-04 | 1.36E-03 | 4.14E-04 | 0.00E+00 | 0.00E+00 | 0.4359 |
| Capparaceae | *Capparis fascicularis* | 1139.37 | | 0.0744 | 0.1245 | 0.0057 | 0.0194 | 0.0598 | 0.0099 | 0.0867 | 1.46E-02 | 8.99E-03 | 0.00E+00 | 9.46E-05 | 5.68E-04 | 2.93E-03 | 1.14E-03 | 9.46E-05 | 0.00E+00 | 0.4088 |
| Capparaceae | *Capparaceae sp* | 611.25 | 9 | 0.0661 | 0.1331 | 0.0063 | 0.0194 | 0.0712 | 0.0094 | 0.0887 | 1.28E-02 | 9.71E-03 | 9.16E-05 | 1.83E-04 | 3.66E-04 | 1.92E-03 | 1.92E-03 | 2.75E-04 | 0.00E+00 | 0.4215 |
| Cariaceae | *Carica papaya* | 372 | | 0.0588 | 0.3111 | 0.0048 | 0.0052 | 0.0104 | 0.0166 | 0.0131 | 2.39E-03 | 7.02E-05 | 1.40E-05 | 2.81E-05 | 0.00E+00 | 2.39E-04 | 0.00E+00 | 0.00E+00 | 0.00E+00 | 0.4228 |
| Cleomaceae | *Cleome africana* | 484.11 | | 0.0735 | 0.2358 | 0.0122 | 0.0221 | 0.1306 | 0.0088 | 0.1201 | 1.58E-02 | 1.49E-02 | 9.97E-05 | 1.99E-04 | 0.00E+00 | 4.99E-04 | 6.98E-04 | 1.99E-04 | 0.00E+00 | 0.6354 |
| Cleomaceae | *Coalisina angustifolia* | 342.3 | 10 | 0.1041 | 0.1621 | 0.0076 | 0.0133 | 0.0747 | 0.0062 | 0.0982 | 1.07E-02 | 7.96E-04 | 1.33E-04 | 3.98E-04 | 1.33E-04 | 0.00E+00 | 0.00E+00 | 0.00E+00 | 0.00E+00 | 0.4783 |
| Cleomaceae | *Cleome arabica* | 464.55 | | 0.0757 | 0.2393 | 0.0121 | 0.022 | 0.1364 | 0.0082 | 0.1171 | 1.42E-02 | 1.38E-02 | 0.00E+00 | 3.89E-04 | 0.00E+00 | 9.72E-05 | 9.72E-05 | 9.72E-05 | 0.00E+00 | 0.6394 |
| Cleomaceae | *Polanisia sp* | 709.05 | | 0.2838 | 0.1974 | 0.0087 | 0.0212 | 0.0746 | 0.0045 | 0.1143 | 1.13E-02 | 3.64E-04 | 0.00E+00 | 9.10E-05 | 0.00E+00 | 9.10E-05 | 9.10E-05 | 0.00E+00 | 0.00E+00 | 0.7165 |
| Cleomaceae | *Coalisina paradoxa* | 484.11 | | 0.1414 | 0.1237 | 0.0097 | 0.0341 | 0.0859 | 0.0084 | 0.1387 | 2.76E-02 | 9.02E-04 | 8.20E-05 | 1.64E-04 | 0.00E+00 | 4.10E-04 | 4.10E-04 | 0.00E+00 | 0.00E+00 | 0.5715 |

| | | | | | | | | | | | | | | | | | | | | |
|---|---|---|---|---|---|---|---|---|---|---|---|---|---|---|---|---|---|---|---|---|
| Cleomaceae | *Cleomaceae sp* | 322.74 | | 0.2117 | 0.139 | 0.0083 | 0.0284 | 0.0647 | 0.0064 | 0.11 | 1.80E-02 | 3.70E-04 | 2.22E-04 | 7.40E-05 | 0.00E+00 | 4.44E-04 | 1.63E-03 | 0.00E+00 | 7.40E-05 | 0.5891 |
| Cleomaceae | *Cleome violacea* | 474.33 | | 0.1074 | 0.1905 | 0.0094 | 0.021 | 0.1124 | 0.0045 | 0.1136 | 1.34E-02 | 1.35E-02 | 1.09E-04 | 5.45E-04 | 0.00E+00 | 3.27E-04 | 6.54E-04 | 0.00E+00 | 0.00E+00 | 0.5874 |
| Cleomaceae | *Melidiscus giganteus* | 420.54 | 10 | 0.1345 | 0.1361 | 0.0094 | 0.0381 | 0.0889 | 0.0097 | 0.1485 | 2.26E-02 | 1.58E-03 | 7.55E-05 | 2.26E-04 | 0.00E+00 | 7.55E-04 | 4.53E-04 | 0.00E+00 | 0.00E+00 | 0.5909 |
| Cleomaceae | *Arivela viscosa* | 268.95 | 16,17 | 0.0521 | 0.1302 | 0.0099 | 0.0231 | 0.0931 | 0.0067 | 0.1282 | 1.94E-02 | 2.19E-03 | 0.00E+00 | 1.29E-04 | 0.00E+00 | 1.29E-04 | 2.58E-04 | 0.00E+00 | 0.00E+00 | 0.4654 |
| Cleomaceae | *Cleomella serrulata* | NA | | NA | NA | NA | NA | NA | NA | NA | NA | NA | NA | NA | NA | NA | NA | NA | NA | NA |
| Cleomaceae | *Gynanadropsis gynandra* | 1124.7 | 17,18 | 0.2219 | 0.2522 | 0.0055 | 0.0183 | 0.0476 | 0.0049 | 0.0756 | 1.29E-02 | 8.43E-04 | 0.00E+00 | 1.40E-04 | 0.00E+00 | 2.81E-04 | 7.02E-04 | 0.00E+00 | 0.00E+00 | 0.6408 |
| Cleomaceae | *Polanisia dodecandra* | 733.5 | 10 | 0.3409 | 0.1883 | 0.0072 | 0.0154 | 0.0596 | 0.0042 | 0.0945 | 1.31E-02 | 1.68E-04 | 1.68E-04 | 0.00E+00 | 0.00E+00 | 1.68E-04 | 8.42E-05 | 0.00E+00 | 0.00E+00 | 0.7239 |
| Cleomaceae | *Polanisia graveolens* | 723.72 | | 0.3499 | 0.1821 | 0.0078 | 0.0181 | 0.054 | 0.0042 | 0.096 | 1.20E-02 | 2.56E-04 | 8.52E-05 | 2.56E-04 | 0.00E+00 | 8.52E-05 | 8.52E-05 | 0.00E+00 | 0.00E+00 | 0.7248 |
| Cleomaceae | *Polanisia trachysperma* | 699.27 | 10 | 0.288 | 0.2218 | 0.0071 | 0.0198 | 0.0657 | 0.0051 | 0.1022 | 1.18E-02 | 6.44E-04 | 0.00E+00 | 9.19E-05 | 0.00E+00 | 9.19E-05 | 2.76E-04 | 0.00E+00 | 0.00E+00 | 0.7224 |
| Cleomaceae | *Tarenaya hassleriana* | 435.21 | 10 | 0.2576 | 0.1102 | 0.0069 | 0.0221 | 0.046 | 0.0052 | 0.0855 | 1.39E-02 | 1.51E-03 | 6.57E-05 | 1.31E-04 | 0.00E+00 | 1.97E-04 | 5.19E-03 | 0.00E+00 | 0.00E+00 | 0.5544 |
| Moringaceae | *Moringa oleifera* | 386.31 | 11 | 0.0596 | 0.1354 | 0.0147 | 0.0241 | 0.0694 | 0.0134 | 0.0972 | 1.86E-02 | 6.46E-04 | 0.00E+00 | 3.23E-04 | 0.00E+00 | 4.84E-04 | 1.61E-04 | 0.00E+00 | 0.00E+00 | 0.4339 |
| Resadaceae | *Ochradenus baccatus* | 870.42 | 24,28 | 0.2406 | 0.1654 | 0.0017 | 0.0091 | 0.0349 | 0.002 | 0.0424 | 4.27E-03 | 1.86E-04 | 1.86E-04 | 1.86E-04 | 0.00E+00 | 1.86E-04 | 1.86E-04 | 0.00E+00 | 0.00E+00 | 0.5014 |
| Resadaceae | *Reseda ordorata* | 498.78 | | 0.1566 | 0.3133 | 0.0022 | 0.0091 | 0.0562 | 0.0046 | 0.0398 | 9.51E-03 | 4.42E-04 | 4.42E-04 | 4.42E-04 | 0.00E+00 | 0.00E+00 | 0.00E+00 | 0.00E+00 | 0.00E+00 | 0.5926 |

**Supplemental Table 2.** List of mitochondrial and chloroplast sequences used for data filtering.

| Mitochondrial reads | Chloroplast reads |
|---|---|
| Brassica napus mitochondrial linear plasmid, complete sequence | Arabidopsis thaliana chloroplast, complete genome |
| Brassica napus mitochondrion, complete genome | Aethionema cordifolium chloroplast, complete genome |
| Carica papaya mitochondrion, complete genome | Aethionema grandiflorum chloroplast, complete genome |
| Brassica oleracea mitochondrion, complete genome | Olimarabidopsis pumila chloroplast, complete genome |
| Brassica carinata mitochondrion, complete genome | Arabis hirsuta chloroplast, complete genome |
| Brassica juncea mitochondrion, complete genome | Barbarea verna chloroplast, complete genome |
| Brassica rapa subsp. campestris mitochondrion, complete genome | Capsella bursa-pastoris chloroplast, complete genome |
| Raphanus sativus mitochondrion, complete genome | Crucihimalaya wallichii chloroplast, complete genome |
| Batis maritima mitochondrion, complete genome | Draba nemorosa chloroplast, complete genome |
| Brassica nigra mitochondrion, complete genome | Lepidium virginicum chloroplast, complete genome |
| Sinapis arvensis mitochondrion, complete genome | Lobularia maritima chloroplast, complete genome |
| Arabis alpina mitochondrion, complete genome | Nasturtium officinale chloroplast, complete genome |
| Arabidopsis thaliana ecotype Col-0 mitochondrion, complete genome | Carica papaya chloroplast, complete genome |
| Boechera stricta mitochondrion, complete genome | Brassica rapa subsp. pekinensis chloroplast, complete genome |
| | Brassica napus chloroplast, complete genome |
| | Pachycladon enysii chloroplast, complete genome |
| | Pachycladon cheesemanii chloroplast, complete genome |
| | Arabis alpina complete chloroplast genome |
| | Raphanus sativus chloroplast, complete genome |
| | Capsella rubella chloroplast, complete genome |
| | Eutrema salsugineum chloroplast, complete genome |
| | Brassica juncea chloroplast, complete genome |
| | Isatis tinctoria voucher cp-824 chloroplast, complete genome |
| | Capsella grandiflora chloroplast, complete genome |
| | Schrenkiella parvula chloroplast, complete genome |
| | Eutrema yunnanense chloroplast, complete genome |
| | Eutrema heterophyllum chloroplast, complete genome |
| | Cochlearia borzaeana chloroplast genome, complete sequence, isolate Cbor_1063 |
| | Cochlearia islandica chloroplast genome, complete sequence, isolate Cisla_1233 |
| | Cochlearia pyrenaica chloroplast genome, complete sequence, isolate Cpyr_0260 |
| | Cochlearia tridactylites chloroplast genome, complete sequence, isolate Ctri_1288 |
| | Ionopsidium acaule chloroplast genome, complete sequence, isolate Iacau_1072 |
| | Arabidopsis arenosa chloroplast genome, complete sequence, specimen voucher PUSTE-A |
| | Arabidopsis cebennensis chloroplast genome, complete sequence, specimen voucher 138R-17 |
| | Arabidopsis pedemontana chloroplast genome, complete sequence, specimen voucher 114R-03 |
| | Camelina sativa chloroplast genome, complete sequence, specimen voucher B-2007-0434-12 |
| | Eutrema halophilum chloroplast, complete genome |
| | Eutrema botschantzevii chloroplast, complete genome |
| | Arabidopsis arenicola chloroplast genome, complete sequence, isolate 2329 |
| | Arabidopsis croatica chloroplast genome, complete sequence, isolate ACR |

| | |
|---|---|
| | Arabidopsis neglecta chloroplast genome, complete sequence, isolate AA095-F |
| | Arabidopsis petrogena chloroplast genome, complete sequence, isolate SK08-h |
| | Arabidopsis suecica chloroplast genome, complete sequence, isolate AS459 |
| | Arabidopsis umezawana chloroplast genome, complete sequence, isolate 504865 |
| | Brassica nigra chloroplast, complete genome |
| | Pugionium dolabratum chloroplast, complete genome |
| | Pugionium cornutum chloroplast, complete genome |
| | Cakile arabica chloroplast, complete genome |
| | Orychophragmus diffusus chloroplast, complete genome |
| | Orychophragmus taibaiensis chloroplast, complete genome |
| | Orychophragmus hupehensis chloroplast, complete genome |
| | Alliaria grandifolia chloroplast, complete genome |
| | Cardamine limprichtiana chloroplast, complete genome |
| | Alyssum desertorum voucher ADES20160707 chloroplast, complete genome |
| | Megadenia pygmaea chloroplast, complete genome |
| | Matthiola incana chloroplast, complete genome |
| | Solms-laubachia eurycarpa chloroplast, complete genome |
| | Megacarpaea delavayi chloroplast, complete genome |
| | Neotorularia korolkowii chloroplast, complete genome |
| | Thlaspi arvense chloroplast, complete genome |
| | Lepidium meyenii chloroplast, complete genome |
| | Tarenaya hassleriana chloroplast, complete genome |
| | Arabidopsis lyrata chloroplast, complete genome |
| | Arabidopsis halleri chloroplast, complete genome |
| | Aethionema arabicum chloroplast, complete genome |
| | Arabidopsis lyrata subsp. lyrata chloroplast DNA, complete genome, strain: MN47 |
| | Sinapis arvensis chloroplast, complete genome |
| | Hesperis matronalis chloroplast, complete genome |
| | Hesperis sylvestris chloroplast, complete genome |
| | Lobularia libyca chloroplast, complete genome |
| | Morettia canescens chloroplast, complete genome |
| | Braya humilis chloroplast, complete genome |
| | Bunias erucago chloroplast genome, complete sequence, specimen voucher OSBU:12478 |
| | Bunias orientalis chloroplast genome, complete sequence, specimen voucher RO12 |
| | Arabis flagellosa Kifune chloroplast DNA, complete genome |
| | Bretschneidera sinensis chloroplast, complete genome |
| | Draba oreades chloroplast, complete genome |
| | Sisymbrium irio chloroplast J04 DNA, complete genome |
| | Biscutella baetica isolate Bba1 chloroplast, complete genome |
| | Biscutella lyrata isolate Bly2 chloroplast, complete genome |
| | Heldreichia bupleurifolia isolate Hbu3 chloroplast, complete genome |
| | Lunaria rediviva isolate Lre4 chloroplast, complete genome |
| | Ricotia aucheri isolate Rau5 chloroplast, complete genome |

| | |
|---|---|
| | Ricotia carnosula isolate Rca6 chloroplast, complete genome |
| | Ricotia cretica isolate Rcr7 chloroplast, complete genome |
| | Ricotia davisiana isolate Rda8 chloroplast, complete genome |
| | Ricotia isatoides isolate Ris9 chloroplast, complete genome |
| | Ricotia lunaria isolate Rlu10 chloroplast, complete genome |
| | Brassica rapa isolate c13 chloroplast, complete genome |
| | Brassica oleracea isolate HDEM chloroplast, complete genome |
| | Moringa oleifera chloroplast, complete genome |

**Supplemental Table 3.** Data processing summary.

| Family | Species | Genome_Size_Mbp | Raw_reads | Filtered_reads | Total_reads_sampled | Total_reads_clustered | Percent_unannotated |
|---|---|---|---|---|---|---|---|
| Brassicaceae | *Aethionema arabicum* | 933.99 | 17,515,712 | 13,704,622 | 500,000 | 298,181 | 0.787799994 |
| Brassicaceae | *Meniocus linifolius* | 899.76 | 19,025,676 | 17,900,990 | 500,000 | 205,355 | 0.866199996 |
| Brassicaceae | *Barbarea bracteosa* | 268.95 | 17,958,660 | 16,274,282 | 500,000 | 270,845 | 1.117800003 |
| Bataceae | *Batis maritima* | 356.97 | 16,618,110 | 15,093,764 | 500,000 | 177,780 | 0.171599998 |
| Capparaceae | *Boscia sp* | 410.76 | 21,741,004 | 19,951,552 | 500,000 | 195,140 | 1.424399998 |
| Capparaceae | *Capparaceae sp* | 259.17 | 18,732,834 | 15,702,982 | 500,000 | 217,968 | 1.441000001 |
| Brassicaceae | *Cakile maritima* | 1383.87 | 20,109,922 | 17,345,778 | 500,000 | 290,826 | 0.769600012 |
| Brassicaceae | *Calepina irregularis* | 254.28 | 18,945,186 | 14,996,774 | 500,000 | 230,865 | 0.667199996 |
| Capparaceae | *Capparis fascicularis* | 1139.37 | 20,757,542 | 19,626,330 | 500,000 | 204,412 | 0.881599999 |
| Capparaceae | *Capparis tomentosa* | 611.25 | 18,850,070 | 16,916,978 | 500,000 | 210,754 | 0.937799997 |
| Brassicaceae | *Capsella bursa-pastoris* | 420.54 | 19,924,400 | 17,596,938 | 500,000 | 244,673 | 1.018199992 |
| Brassicaceae | *Cardamine hirsuta* | 489 | 14,327,238 | 13,353,298 | 500,000 | 236,564 | 0.893800004 |
| Cariaceae | *Carica papaya* | 372 | 26,670,784 | 22,740,686 | 500,000 | 211,407 | 2.305599995 |
| Brassicaceae | *Caulanthus amplexicaulis* | 484.11 | 17,276,330 | 13,566,798 | 500,000 | 227,219 | 0.618400004 |
| Brassicaceae | *Chorispora tenella* | 366.75 | 21,426,658 | 17,199,758 | 500,000 | 255,923 | 0.669999999 |
| Cleomaceae | *Cleome africana* | 484.11 | 14,535,678 | 12,523,252 | 500,000 | 317,696 | 1.167200004 |
| Cleomaceae | *Coalisina angustifolia* | 342.3 | 17,685,930 | 16,598,982 | 300,000 | 143,503 | 1.172000003 |
| Cleomaceae | *Cleome arabica* | 464.55 | 15,921,148 | 12,608,460 | 500,000 | 319,725 | 1.200000004 |
| Cleomaceae | *Polanisia sp* | 709.05 | 16,484,740 | 14,702,180 | 500,000 | 358,263 | 0.9018 |
| Cleomaceae | *Coalisina paradoxa* | 484.11 | 20,093,758 | 17,333,978 | 500,000 | 285,751 | 1.307799997 |
| Cleomaceae | *Cleomaceae sp* | 322.74 | 20,481,152 | 17,552,776 | 500,000 | 294,564 | 1.399400005 |
| Cleomaceae | *Cleome violacea* | 474.33 | 15,366,960 | 13,381,026 | 400,000 | 234,951 | 1.143750005 |
| Cleomaceae | *Melidiscus giganteus* | 420.54 | 18,036,600 | 15,768,938 | 500,000 | 295,440 | 1.648799998 |
| Cleomaceae | *Arivela viscosa* | 268.95 | 18,197,094 | 14,690,618 | 400,000 | 186,177 | 2.092499998 |
| Brassicaceae | *Cochlearia offcinalis* | 734 | 19,571,568 | 17,680,152 | 500,000 | 309,075 | 0.633400004 |
| Brassicaceae | *Crambe hispanica* | 660.15 | 17,962,804 | 16,030,190 | 500,000 | 237,458 | 0.847600007 |
| Brassicaceae | *Descurainia pinnata* | 371.64 | 21,064,170 | 18,911,684 | 500,000 | 246,938 | 0.555600003 |
| Brassicaceae | *Descurainia sophioides* | 195.6 | 14,299,738 | 11,722,246 | 500,000 | 248,632 | 0.685599997 |
| Brassicaceae | *Diptychocarpus strictus* | 557.46 | 14,976,410 | 13,219,700 | 500,000 | 255,043 | 0.805800004 |
| Brassicaceae | *Eruca vesicaria* | 709.05 | 19,393,468 | 16,238,972 | 500,000 | 267,256 | 0.8802 |
| Brassicaceae | *Erysimum cheiranthoides* | 420.54 | 19,255,900 | 14,718,730 | 500,000 | 258,177 | 1.210200004 |
| Brassicaceae | *Euclidium syriacum* | 298.29 | 16,998,370 | 15,169,108 | 500,000 | 223,832 | 0.641799998 |
| Brassicaceae | *Farsetia aegyptica* | 2004.9 | 17,029,968 | 16,121,632 | 500,000 | 239,500 | 0.934600003 |
| Brassicaceae | *Goldbachia laevigata* | 547.68 | 19,699,580 | 18,035,220 | 500,000 | 231,487 | 0.6218 |
| Brassicaceae | *Guillenia lasiophylla* | 621.03 | 11,213,854 | 9,994,886 | 500,000 | 189,388 | 1.013799997 |
| Cleomaceae | *Gynanadropsis gynandra* | 1124.7 | 18,472,870 | 16,563,780 | 500,000 | 320,397 | 0.499200002 |
| Brassicaceae | *Hesperis matronalis* | 3261.63 | 15,614,640 | 15,174,086 | 500,000 | 333,859 | 0.537600005 |
| Brassicaceae | *Hirschefeldia incana* | 552.57 | 23,055,074 | 21,141,192 | 500,000 | 232,731 | 1.167399996 |
| Brassicaceae | *Iberis amara* | 831.3 | 13,691,982 | 13,178,566 | 500,000 | 270,980 | 1.225400004 |
| Brassicaceae | *Isatis lusitanica* | 381.42 | 23,390,698 | 17,191,612 | 500,000 | 282,703 | 0.580399999 |
| Brassicaceae | *Isatis tinctoria* | 616.14 | 18,892,040 | 15,557,688 | 500,000 | 246,272 | 0.846399996 |

| | | | | | | | |
|---|---|---|---|---|---|---|---|
| Brassicaceae | *Lepidium ruderale* | 371.64 | 18,919,410 | 14,656,664 | 500,000 | 195,884 | 0.829600002 |
| Brassicaceae | *Lepidium sativuum* | 591.69 | 18,902,354 | 16,242,402 | 500,000 | 245,618 | 0.521800004 |
| Brassicaceae | *Lobularia maritima* | 312.96 | 19,156,394 | 14,101,930 | 500,000 | 257,902 | 0.571799998 |
| Brassicaceae | *Lunaria annua* | 498.78 | 14,633,200 | 12,998,044 | 500,000 | 313,439 | 0.6024 |
| Brassicaceae | *Malcomia maritima* | 312.96 | 15,816,038 | 13,287,250 | 250,000 | 122,050 | 1.107999997 |
| Brassicaceae | *Matthiola longipetala* | 1599.03 | 20,089,766 | 19,078,766 | 500,000 | 344,981 | 0.498000005 |
| Moringaceae | *Moringa oleifera* | 386.31 | 20,478,894 | 17,095,412 | 500,000 | 216,954 | 0.590999995 |
| Brassicaceae | *Myagrum perfoliatum* | 352.08 | 13,275,434 | 10,829,146 | 400,000 | 220,796 | 0.700749995 |
| Brassicaceae | *Nasturtium officinale* | 430.32 | 16,369,294 | 13,096,042 | 500,000 | 270,006 | 0.548799996 |
| Resadaceae | *Ochradenus baccatus* | 870.42 | 18,680,706 | 17,328,066 | 500,000 | 250,676 | 0.341200004 |
| Brassicaceae | *Olimoarabidopsis pumila* | 410.76 | 14,092,992 | 11,158,530 | 500,000 | 221,195 | 1.072000003 |
| Brassicaceae | *Physaria acutifolia* | 2322.75 | 18,571,300 | 17,404,712 | 500,000 | 361,507 | 1.074200005 |
| Cleomaceae | *Polanisia dodecandra* | 733.5 | 17,592,446 | 16,046,308 | 500,000 | 361,926 | 0.840400003 |
| Cleomaceae | *Polanisia graveolens* | 723.72 | 20,436,352 | 18,930,976 | 500,000 | 362,412 | 0.833199998 |
| Cleomaceae | *Polanisia trachysperma* | 699.27 | 18,164,388 | 14,896,462 | 500,000 | 361,223 | 0.827400004 |
| Brassicaceae | *Sisymbrium leucocladum* | 498.78 | 21,306,942 | 17,347,804 | 500,000 | 284,673 | 0.703999996 |
| Brassicaceae | *Psychine stylosa* | 518.34 | 18,672,344 | 15,813,984 | 500,000 | 246,922 | 0.815000003 |
| Resadaceae | *Reseda ordorata* | 498.78 | 20,059,938 | 17,972,876 | 500,000 | 296,311 | 0.307800004 |
| Brassicaceae | *Rorippa islandica* | 567.24 | 14,323,126 | 12,382,424 | 500,000 | 244,939 | 1.065999997 |
| Brassicaceae | *Schizopetalum walkeri* | 1775.07 | 19,473,154 | 18,463,848 | 500,000 | 317,020 | 0.339599997 |
| Brassicaceae | *Sinapis alba* | 557.46 | 19,804,700 | 16,543,166 | 500,000 | 249,449 | 0.833399989 |
| Brassicaceae | *Sisymbrium brassiciforme* | 371.64 | 18,157,024 | 15,596,368 | 500,000 | 247,748 | 0.779399995 |
| Brassicaceae | *Brassicaceae sp* | 1036.68 | 14,359,594 | 12,039,944 | 500,000 | 258,811 | 0.911999998 |
| Brassicaceae | *Streptanthus arizonicus* | 1090.47 | 20,032,530 | 17,795,582 | 500,000 | 291,308 | 0.391000003 |
| Brassicaceae | *Streptanthus heterophyllys* | 748.17 | 20,440,916 | 18,181,630 | 500,000 | 189,327 | 0.588599995 |
| Cleomaceae | *Tarenaya hassleriana* | 435.21 | 24,418,660 | 22,321,972 | 500,000 | 277,209 | 1.194800003 |
| Brassicaceae | *Teesdalia nudicaulis* | 557.46 | 17,158,882 | 15,882,064 | 500,000 | 277,737 | 0.503399998 |
| Brassicaceae | *Thlaspi arvense* | 205.38 | 13,742,780 | 12,034,208 | 500,000 | 326,494 | 0.530000001 |
| Brassicaceae | *Turritis glabra* | 919.32 | 21,721,584 | 19,358,608 | 500,000 | 230,979 | 1.205600012 |
| Cleomaceae | *Cleomella serrulata* | N/A | N/A | N/A | N/A | N/A | N/A |

**Supplemental Table 4.** OUwie weighted AICc scores. Tests conducted are specified below as selection regime ~ modelled phenotype. BM1; single-rate Brownian motion, BMS; Brownian motion with different rate parameters for each state on a tree, OU1; Ornstein-Uhlenbeck model with a single optimum for all species, OUM; Ornstein-Uhlenbeck model with different state means and a single alpha and sigma^2 acting all selective regimes, OUMV; Ornstein-Uhlenbeck model that assumes different state means as well as multiple sigma^2, OUMA; Ornstein-Uhlenbeck model that assumes different state means as well as multiple alpha, OUMVA; Ornstein-Uhlenbeck model that assumes different state means as well as multiple alpha and sigma^2 per selective regime. Asterisk (*) indicated analyses which returned the warning, "You might not have enough data to fit this model well".

| Simmap | Test Conducted | OUM | OU1 | BM1 | BMS | OUMV | OUMA | OUMVA |
|---|---|---|---|---|---|---|---|---|
| 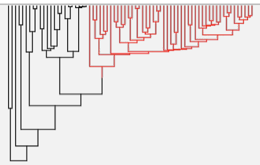 | AtAlpha/All ~ Total TE | 0.03478328 | 0.10264773 | 0.059294 | ***0.4075957*** | 0.05291083 | 0.10698661 | 0.23578184 |
| | AtAlpha/All ~ Gyspy | 0.23758937 | ***0.3186865*** | 0.00295855 | 0.00190546 | 0.14597727 | 0.20500947 | 0.08787339 |
| | AtAlpha/All ~ Copia | 0.29076304 | ***0.3509629*** | 0.00001552 | 0.000009 | 0.15800576 | 0.15221466 | 0.0480291 |
| 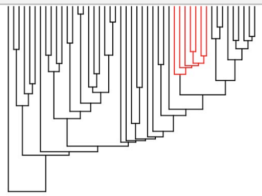 | Brassiceae/Brassicaceae ~ Total TE | 0.09524774 | ***0.2741048*** | 0.0795574 | 0.08170141 | 0.21402713* | 0.15597822* | 0.09938331* |
| | Brassiceae/Brassicaceae ~ Gyspy | 0.11124287 | ***0.3679953*** | 0.32010442 | 0.10217491 | 0.04138997* | 0.03731004* | 0.0197825* |
| | Brassiceae/Brassicaceae ~ Copia | 0.18430769 | ***0.5635547*** | 0.01588842 | 0.00555477 | 0.10066517* | 0.09134368* | 0.03868553* |
| 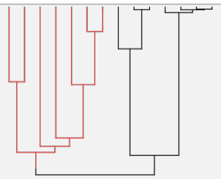 | ThAlpha/Cleomaceae ~ Total RE | 0.09367094* | 0.1104097* | ***0.5775217**** | 0.19361534* | 0.01500279* | 0.00897249* | 0.00080708* |
| | ThAlpha/Cleomaceae ~ Gyspy | 0.27132915* | ***0.3576261**** | 0.12686794* | 0.11919859* | 0.05904368* | 0.06326126* | 0.00267328* |
| | ThAlpha/Cleomaceae ~ Copia | 0.05613062* | ***0.3893679**** | 0.17259922* | 0.36403269* | 0.00970944* | 0.00778289* | 0.00037727* |
| 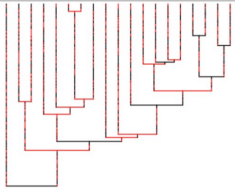 | Neo/BMAP ~ Total RE | 0.09177786* | 0.1752389* | ***0.4870293**** | 0.16334972* | 0.01504765* | 0.05696647* | 0.01059003* |
| | Neo/BMAP ~ Gyspy | 0.10288377* | 0.1438035* | ***0.4304128**** | 0.15543752* | 0.01833382* | 0.14525589* | 0.00387266* |
| | Neo/BMAP ~ Copia | 0.37400051* | ***0.4134385**** | 0.01666794* | 0.00857456* | 0.11452479* | 0.06447625* | 0.00831744* |
